# Structure-guided optimization of light-activated chimeric G-protein coupled receptors

**DOI:** 10.1101/2021.12.13.472518

**Authors:** Alexandra-Madelaine Tichy, Wang Lok So, Elliot J. Gerrard, Harald Janovjak

## Abstract

G-protein coupled receptors (GPCRs) are the largest human receptor family and involved in virtually every physiological process. One hallmark of GPCR function is the specific coupling of activated receptors to selected downstream signaling pathways. The ability to tune this coupling would permit the development of receptors with new capabilities. GPCRs and G-proteins have been recently resolved structurally at high resolution, but this information was in only very few cases harnessed for a rational engineering of these protein complexes. Here, we demonstrate the structure-guided optimization of coupling in chimeric light-activated GPCRs (OptoXRs). Our hypothesis was that the incorporation of structural GPCR-Gα contacts will lead to improved receptor activity. We first evaluated structure-based alignments as complements to existing sequence-based methods for generation of chimeric receptors. We then show in a prototypical light-activated β_2_AR that inclusion of α-helical residues forming structural contacts to Gα resulted in receptors with 7- to 20-fold increased function compared to other design strategies. In turn, elimination of GPCR-Gα contacts diminished function. Finally, the efficient receptor design served as a platform for the optimization of a further light-activated receptor and spectral tuning of the photoreceptor core domain. Our work exemplifies how increased OptoXR potency and new functionalities can be achieved through structure-based design towards targeted inputs into cells and cellular networks.

## INTRODUCTION

G-protein coupled receptors (GPCRs) comprise the largest family of membrane receptors in the human genome. They are expressed broadly and regulate key biological processes, ranging from organism development to metabolism and brain function^1,2^. In line with their abundance and physiological importance, GPCRs are also exceptional therapeutic targets with ~30% of all prescription drugs acting on members of this receptor superfamily^3,4^.

Canonical signaling downstream of GPCRs is mediated through coupling to heterotrimeric G-proteins, which are often classified into four main groups according to their Gα subunits (the Gα_s_, Gα_i/o_, Gα_q_ and Gα_12/13_ groups). Gα subunits principally act on adenylyl cyclase to increase (Gα_s_) or decrease (Gα_i/o_) cAMP levels, on phospholipase C (PLC) to alter Ca^2+^ signaling (Gα_q_) or on Rho GTPases (Gα_12/13_) to modulate motor proteins and the cytoskeleton (**Fig. 1a**). In addition, associated Gβ and Gγ subunits activate ion channels and a wide range of enzymes upon dissociation of the trimeric complex. In humans, at least 16 Gα subunits, 6 Gβ subunits and 13 Gγ subunits have been identified, which can assemble in a multitude of combinations leading to functionally-distinct outcomes^5–7^. GPCRs can also signal *via* β-arrestin scaffolding proteins ultimately to diverse signaling pathways (**Fig. 1a**). Importantly, the expression of multiple GPCRs and G-proteins in many cell types results in combinatorial complexity and dictates that specific GPCR-G-protein interactions guide signaling outcomes^5,8^.

**Fig. 1:**
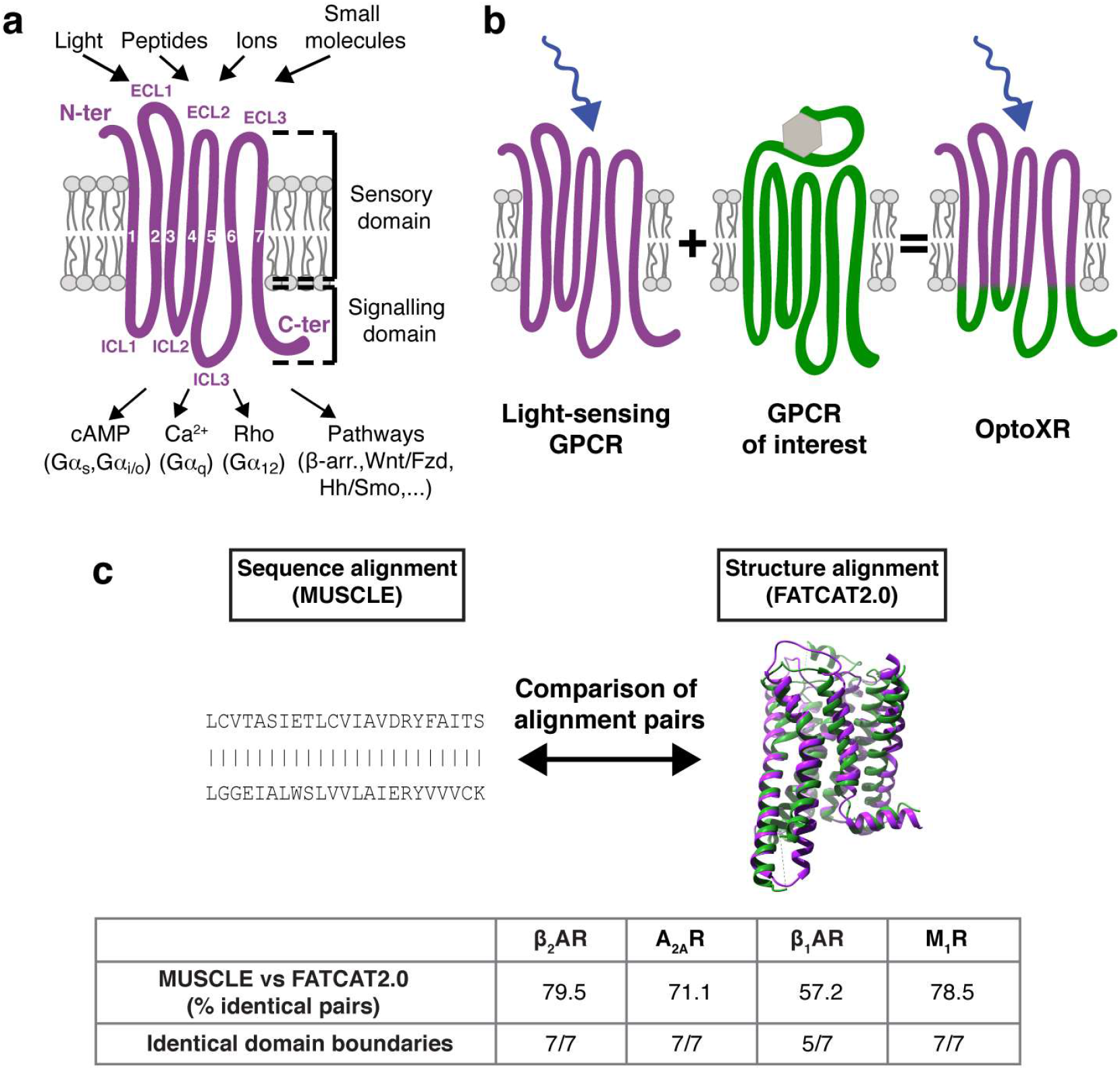
Functional principle of OptoXRs and alignment methodologies. **(a)** GPCRs sense a variety of external stimuli using their extracellular and TM domains. Downstream signaling is mediated by residues that face the intracellular environment. **(b)** In chimeric OptoXRs, the signaling elements of a target GPCR of interest is fused to the sensory elements of a light-activated GPCR. **(c)** Top: Sequence- and structure-based alignments were compared. Bottom: Quantification of alignment similarity expressed either as the percentage of identical residue pairs (top row; see **Supplementary Fig. S1** for a definition of residue pairs) or as the number of identical domain boundaries (bottom row; see **Supplementary Fig. S2** for a definition of domain boundaries).

Consequences of GPCR signaling are most commonly studied using pharmacological (e.g., agonists, antagonists or allosteric modulators) or genetic methods (e.g., receptor overexpression or knock-out) in cells and animals. As a complement, approaches have been developed to (in)activate GPCR signaling on shorter time scales and finer spatial scales than those dictated by drug diffusion or pharmacokinetics and those associated with genetic alterations^9,10^. One such approach are light-activated GPCRs, which are either repurposed from naturally light-sensing mammalian tissues^11–13^ or from invertebrates and fish^14–17^, or are engineered as chimeric GPCRs (termed OptoXRs)^18,19^. Albeit of different sources, these light-activated GPCRs all permit spatio-temporal activation of signaling pathways by light in the emerging field of optogenetics^20,21^.

The purpose-built OptoXRs take advantage of conserved structure-function relationships by substituting the downstream coupling domains of a light-gated GPCR, such as the mammalian dim light sensor rhodopsin (rho), with those of a naturally ligand-gated target GPCR (**Fig. 1b**)^18,19^. The desired engineering outcome is that the chimeric GPCR can be activated by light but retains the signaling properties of the target receptor. OptoXRs have been expressed in neuronal and non-neuronal cell types *ex vivo* and *in vivo* using viral delivery or in transgenic animals and activated with light from a variety of sources^22–24^. Light control was often achieved *in vivo* without supplementation of the *cis*-isoform retinal cofactor^18,25,26^, but supplementation protocols have also been developed^22^. The OptoXR methodology has been applied to a number of target GPCRs by us and others (see **Supplementary Table S1** for a -to our best knowledge-comprehensive list of OptoXRs). Notably, the design principle underlying most published receptors (see column “Domain Boundaries” in **Supplementary Table S1**) was derived from a pioneering study^19^ that employed a trial-and-error screening strategy (i.e., various receptor residues and receptor domains were swapped and the emerging receptors tested functionally). This original and more recent trial-and-error designs have put forward the notion that intracellular loops (ICLs) are dominant determinants of function in OptoXRs^19,27^, but this has never been tested systematically^9^. The increasing application of chimeric and natural light-activated GPCRs not only as basic research tools but even potential future treatment strategies^23,28,29^ calls for attempts to rationally engineer OptoXRs to, e.g., improve or diversify their function.

In the past decade, the structural information available for GPCRs has increased dramatically to encompass more than 600 high-resolution structures for more than 150 unique receptors^1,30^. Detailed analyses of these structures have provided insights into GPCR structure-function relationships, such as the universal heptahelical transmembrane (TM) domain, the translation and rotation of specific TM α-helices upon receptor activation or the switch-like behavior of conserved residue networks^31–35^. Following from early successes using X-ray crystallography^36^, it was the advent of cryo-electron microscopy (cryo-EM) that revealed a number of active state and G-protein bound structures for members of several major GPCR classes^37–39^. Albeit structures of GPCR-Gα complexes allow identifying key molecular interactions between GPCRs and their downstream transducers^5,36,40^, this knowledge has been in only very few cases harnessed for the development of receptors with rationally-designed downstream functions^9,41^. Specifically, orthogonal Gα subunits^42^ or GPCRs with promiscuous downstream coupling^43^ were designed using modelling, molecular dynamics simulations and *in silico*/experimental mutagenesis. It remains to be further explored to what extent GPCR function can be redesigned using strategies that build on the identification of GPCR-G-protein contacts. Such strategies may complement seminal work in which downstream coupling in wildtype (WT) and chimeric GPCRs was altered through residue substitutions that were derived from functional screens or biochemical assays^26,44,45^.

Here, we optimized OptoXR function by harnessing high-resolution GPCR-Gα structures. Besides potential future applications in optogenetics, we chose OptoXRs as a model system because an efficient switch in coupling, e.g. from the native Gα_i/o_ preference of rho to a different Gα group of the target receptor, is required for the faithful function of these chimeric receptors. We first explored if atomic structure superpositions are a required complement to sequence alignments for determination of functional domain boundaries. We then developed rational structure analysis and receptor design strategies. Focusing on GPCRs in complex with their restive Gα subunits, we showed that inclusion of TM α-helical residues that form contacts to Gα greatly improved functionality of a light-activated β_2_AR receptor. Finally, we demonstrated optimization of a second receptor and tuning of spectral properties. Collectively, the major advance of this work is a demonstration of structure-guided reengineering of GPCR-Gα coupling yielding improved and diversified OptoXRs.

## METHODS

### Vectors and constructs

OptoXR genes were designed as described in the main text and ordered as synthetic genes (gBlocks, Integrated DNA Technologies). Protein and nucleotide sequences of all engineered receptors can be found in **Supplementary Table S3** and **Supplementary Table S4**, respectively. Genes were amplified in polymerase chain reactions (PCRs) using oligonucleotides with XhoI and KpnI restriction site overhangs and digested with the respective enzymes (NEB). pcDNA3.1 (Thermo Fisher Scientific) was digested with the same enzymes and ligated with the amplified receptors using T4 DNA ligase (Promega).

For β-arrestin recruitment assays, receptors were tagged with Rluc8^46^. Receptor genes were amplified without a stop codon using oligonucleotides with BamHI and XhoI restriction site overhangs and digested with the respective enzymes. A vector containing Rluc8 (a kind gift of P. Sexton, Monash University) was digested with the same enzymes. Receptor genes were ligated into the vector N-terminally of Rluc8 using T4 DNA ligase.

Opto-β_2_AR and Opto-A_2A_R vectors were in the same backbone and described in a previous publication^47^. The Opto-β_2_AR gene encodes for a protein that is identical to those published previously^18,19^. β_2_AR in pcDNA3 was a gift of Robert Lefkowitz (Addgene plasmid #14697)^48^. JellyOp in pcDNA3.1 was a kind gift of Robert Lucas^16^.

Single and double residue substitutions were introduced in Opto-β_2_AR-2.0 using overlapping oligonucleotides (designed in PrimerX (https://www.bioinformatics.org/primerx/) using the QuikChange option) and circular PCRs with a high-fidelity polymerase (Q5, NEB) followed by digestion with DpnI restriction enzyme (NEB).

All genes were verified using Sanger sequencing (MicroMon, Monash University).

### Cell culture

HEK293 cells (Thermo Fisher Scientific; further authenticated by assessing cell morphology and growth rate) were cultured in mycoplasma-free Dulbecco’s modified Eagle medium (DMEM, Gibco) in a humidified incubator with 5% CO_2_ atmosphere at 37°C. Medium was supplemented with 10% FBS, 100 U/ml penicillin and 0.1 mg/ml streptomycin (Gibco). HEK293 cells were reverse transfected using poly(ethyleneimine) (linear, 20 kDA; Polysciences)^49^. Culture medium was changed 6 h after transfection and supplemented with 10 μM 9-*cis*-retinal (Sigma) for experiments with light-activated receptors.

### Real-time cAMP measurements

The real-time cAMP assay employed the GloSensor-22F reporter cloned into pcDNA3.1 as described previously^47^. HEK293 cells were seeded at a density of 5 × 10^4^ cells/well in white 96-well plates and transfected with receptor and reporter at a ratio of 1:1. HEK293 cells endogenously express β_2_AR and were mock transfected with empty vector backbone for β_2_AR conditions. The next day, culture medium was changed to Leibovitz’s L15 (Gibco) starve medium (0.5% FBS, 100 U/ml penicillin and 0.1 mg/ml streptomycin) supplemented with 2 mM D-luciferin (Cayman Chemicals) and 10 μM 9-*cis*-retinal in experiments with light-activated receptors. Cells were incubated for 60 min at room temperature (RT). Luminescence was measured in a microplate reader (Omega, BMG Labtech). Baseline luminescence values were recorded for 8 cycles (90 s/cycle) before stimulation. Cells were stimulated with light or ligand as described in the main text and figure captions. For PTX experiments, cells were incubated with 10 μM PTX (List Labs) overnight (o/n) at 37°C. Light stimulation employed the Xenon white light flash lamp (1 mW nominal output power, 10 ms flash separation) of the microplate reader. The experimental stimulation times scale linearly with the flash number (e.g., 2 s for 200 flashes (fl)). Wavelength was adjusted using optical filters (BMG Labtech; 445 ± 7.5 nm, 480 ± 7.5 nm and 540 ± 7.5 nm; center wavelength (λ) ± full width at half-maximum (FWHM)). Each filter was characterized using a white light source and a spectrophotometer (LI-180, Licor). Xenon lamp light output through the filters was measured using a power meter (LP-1, Sanwa). For the illuminated area of 3 mm^2^, the continuous illumination power density (I) was 1.75, 1.97 and 1.35 mW/cm^2^ for 445, 480 and 540 nm continuously over the stimulation time course.

### Real-time Ca^2+^ measurement

The real-time Ca^2+^ mobilization assay employed mtAequorin in pcDNA3.1. HEK293 cells were seeded at a density of 5 × 10^4^ cells/well in white 96-well plates and transfected with receptor and reporter at a ratio of 1:1. The next day, culture medium was replaced with Hank’s buffered salt solution (HBSS, Sigma) supplemented with 10 μM coelenterazine h (Nanolight) and 10 μM 9-*cis*-retinal in experiments with light-activated receptors. Cells were then incubated for 2 h at 37°C. Luminescence was measured in the microplate reader. Baseline luminescence values were recorded for 10 cycles (1 s/cycle) before stimulation. Cells were stimulated with light (see above) or ligand at concentrations as described in the main text and figure captions. For YM254890 experiments, cells were incubated with 10 μM YM254890 (AdipoGen Life Sciences) for 2 h at 37°C before measurements.

### Real-time β-arrestin2 recruitment

Receptor mediated β-arrestin2 recruitment was assessed using a real-time BRET assay detecting energy transfer between Rluc8-tagged receptors and a mVenus-tagged β-arrestin2. HEK293 cells were seeded at a density of 5 × 10^4^ cells/well in white 96-well plates and transfected with receptor-Rluc8 and β-arrestin2-mVenus at a ratio of 1:4. Culture medium was changed 6 h after transfection and supplemented with 10 μM 9-*cis*-retinal (Sigma) for experiments with light-activated receptors. The next day, culture medium was replaced with 40 μl HBSS supplemented and 10 μM 9-*cis*-retinal for experiments with light-activated receptors. Cells were incubated for 1 h at 37°C. Coelenterazine h was added at a final concentration of 5 μM immediately before measurements. Baseline values were recorded for 8 cycles (90 s/cycle) before stimulation. Cells were stimulated with light (see above) or ligand at concentrations as described in the main text and figure captions. BRET measurements were obtained in the microplate reader using 460 (Rluc8) and 520 nm (mVenus) filters. The BRET ratio was calculated by dividing the emission at 520 nm by the emission at 460 nm.

### Immunofluorescence (IF)

HEK293 cells were seeded on glass coverslips coated with poly-L-ornithine (Sigma) in clear 12-well plates (3 × 10^5^ cells/well) and transfected with 1000 ng receptor plasmid. The next day, under dim red light, cells were washed once with pre-warmed PBS and then fixed with 4% PFA for 10 min at RT. Cells were washed again three times with PBS and blocked with PBS + 0.1% Tween (PBS-T) containing 3% BSA at 37°C for 1 h. Cells were incubated with primary antibody (mouse anti-4D2, 1:500 dilution; Abcam, ab98887) at 4°C o/n, washed three times with PBS, and incubated with secondary antibody (donkey anti-mouse, Alexa488 conjugated, 1:750 dilution; Thermo Fisher Scientific, A21202) at RT for 2 h. Cells were then washed once with PBS, incubated with DAPI (Life Technologies) at a final dilution of 1:4000 at RT for 5 min and washed twice with PBS. Coverslips were mounted on microscopy slides using Mowiol mounting medium (Sigma) and left to dry. Images were acquired using a inverted confocal microscope (C1, Nikon) paired with a Plan Apo VC 60x objective. Cells were imaged at 405 and 488 nm with a scan speed of 21.6 frames per s (caution was taken to avoid oversaturation at both 405 (HV = 74, laser power = 4.2) and 488 nm (HV = 82, laser power = 4.2).

### Flow cytometry (FC)

HEK293 cells were seeded in clear 6-well plates at a density of 1 × 10^6^ cells/well and transfected with 2000 ng receptor plasmid. The next day, under dim red light, cells were washed once with PBS and dispersed using Accutase (Sigma) at 37°C for 15 min. Accutase was neutralized using complete medium before the cell suspension was collected in a 15 ml tube and centrifuged. After removal of supernatant, cells were fixed in 2% PFA at 37°C for 10 min and washed once with PBS. We choose permeabilizing fixation conditions for quantification of total expression levels. Cells were permeabilized with ice-cold methanol on ice for 30 min. Cells were washed twice in FC buffer (PBS supplemented with 2% FBS) and incubated with primary antibody (see above) at RT for 30 min. Cells were then washed once with FC buffer and incubated with secondary antibody (see above) at RT for 30 min. Samples, including mock transfected, unstained and secondary antibody controls, were run on the same day on a LSR Fortessa X-20 Cytometer (BD Biosciences) and data was analyzed using FlowJo software (Flowjo).

### Comparison of sequence and structure alignments

Initial comparisons of sequence alignments and structure alignments were performed on rho (PDB ID: 6CMO) and β_2_AR (PDB ID: 3SN6) using only the receptor chain (chain R). FATCAT2.0 alignments were performed using the pairwise alignment option of the FATCAT2.0 server^50^ (https://fatcat.godziklab.org). Cα atoms were included in the alignments with default alignment parameters. Alignments were manually inspected to determine whether key motifs (E/DRY, CWxP or NPxxY) were correctly paired between the first and the second receptor (e.g., Asp^3×49^ of E/DRY of rho should be paired with Asp^3×49^ of E/DRY of β_2_AR).

Sequence alignments were performed in MUSCLE as previously^47,51,52^ using the sequence of residues that are resolved in the PDB file to enable comparison to structure alignments.

For detailed comparison of FATCAT2.0 and MUSCLE alignments, rho (PDB ID: 6CMO) was also aligned to A_2A_R (PDB ID: 6GDF), β_1_AR (PDB ID: 7JJO) and M_1_R (PDB ID: 6OIJ) followed by quantification of identical pairs in macros written in Igor Pro (Wavemetrics) (**Supplementary Fig. S1**). These MUSCLE and FATCAT2.0 alignments were in a final step manually analyzed to assess whether OptoXR boundaries were at identical positions **Supplementary Fig. S2**).

### GPCR-G-protein interacting residues and visualization

GPCR residues in proximity to the Gα subunit were determined in PyMol as described in the main text and figure captions. If available, multiple receptor structures were analyzed to include all possible binding residues as listed in the figure captions. The obtained molecular contact maps were verified using a second and fully automated software (CMView^53^). PDB IDs of the analyzed structures were as follows for the studied receptors: Rho (6CMO, 6OYA), β_2_AR (3SN6, 7BZ2) and A_2A_R (6GDF). Molecular structure images were generated in PyMol.

### Data analysis and statistics

All pharmacological data were analysed using Prism (9.0, GraphPad). Dose-response data were generated by applying a three-parameter non-linear equation to data extracted from time courses. EC50 values were determined by fitting dose-response data with a sigmoid curve. Statistical measures are provided in the figure captions.

## RESULTS

### Comparison of sequence- and structure-based alignments

The creation of protein chimeras, such as OptoXRs, hinges on the exchange of functionally-equivalent domains between proteins. This is generally achieved by first aligning the proteins of interest, either as primary sequences or as three-dimensional structures, followed by identification of domain boundaries. Engineering of previous OptoXRs exclusively relied on sequence-based alignments, including the prominent chimeras of rho and adrenergic, dopamine, adenosine or muscarinic acetylcholine receptors (**Supplementary Table S1**)^18,47,54^. In a complementary approach, we tested whether alignments of structures would yield chimeric receptors that are different from those generated using sequences. This may indeed be the case due to the additional topological information gained and because sequence similarity between GPCRs from different families can be low (e.g., rho and β_2_AR, A_2A_R, β_1_AR and M_1_R exhibit 20, 21, 22 and 23% sequence identity overall, or 21, 22, 23 and 21% when excluding extra- or intracellular termini, respectively). We performed a quantitative comparison of sequence-based alignments (using MUSCLE as in previous studies^18,47^) and structure-based alignments (using FATCAT2.0^50^). For this analysis, we utilized our own computer code that reports the number of identical residue pairs in two alignments (**Supplementary Fig. S1**). For pairs of rho and β_2_AR, A_2A_R, β_1_AR and M_1_R, we found that sequence- and structure-based alignments differed in ~20 to 40% of residue pairs across the sequence (**Fig. 1c**). To determine if these differences impact the design of OptoXRs, we asked whether alignments were identical at the domain boundaries that were previously utilized to generate these chimeras (**Supplementary Fig. S2a**)^19^. We observed only minimal differences that were limited to one receptor pair (rho:β_1_AR) and single residues (**Fig. 1c**, **Supplementary Fig. S2b**). This result confirms that for these prototypical receptors and these domain boundaries the generation of OptoXR scaffolds is insensitive to the alignment method. This may not necessarily apply to all GPCR pairs as, e.g., long ICLs can potentially confound sequence alignments.

### Receptors designs based on structural GPCR-Gα contacts

After comparison of alignment methods, we went on to redesign a prototypical OptoXR using a structure-guided rationale. We chose β_2_AR as the target receptor as it is well characterized and has been incorporated in a OptoXR previously (the new OptoXR generated here is compared to the existing Opto-β_2_AR in detail below). We closely examined structures of β_2_AR and also of rho in complex with their cognate G-proteins (Gα_s_ for β_2_AR and G_α/t_ for rho)^36,55,56^. Using a distance cutoff of 4 Å, chosen analogously to the first study mapping a GPCR-Gα complex^36^, we determined the receptor residues that are in proximity to Gα (see **Fig. 2a-d** for protein structures, and **Supplementary Fig. S3** and **Supplementary Table S2** for listed contacts). We found that these contacts extend away from the cytosolic ICLs upwards on α-helices TM3, TM5 and TM6 by as much as four α-helical turns (**Fig. 2b,d**). This observation prompted us to test if ILCs are sufficient for GPCR coupling. We engineered a rho receptor containing exclusively the ILCs of β_2_AR (termed Opto-β_2_AR-Loops) and demonstrated expression and membrane localization in HEK293 cells using flow cytometry (FC) and immunofluorescence (IF) (**Fig. 2e,f**) (sequences of all engineered receptors can be found in **Supplementary Tables S3 and S4**). However, upon light stimulation (λ=480 ± 7.5 nm, I=1.97 mW/cm^2^, 40 s, 4000 flashes (fl)) no increase in intracellular cAMP levels was observed, indicating that incorporation of these secondary structure elements in this particular chimeric receptor was not sufficient to confer a downstream signaling ability (**Fig. 2g,h**). We then hypothesized that including an as complete as possible ensemble of structural β_2_AR-Gα_s_ contacts will result in efficient G-protein activation. We therefore designed a variant termed Opto-β_2_AR-All-In where domain boundaries are placed to include all structural contacts (**Supplementary Fig. S3**, **Supplementary Table S2**). This receptor should not only perform efficiently but also provide a basis for examining the contribution of specific contact residues to function (as will be shown below). We first verified expression and localization of Opto-β_2_AR-All-In using FC and IF (**Fig. 2e,f**). We then tested receptor function upon light stimulation (λ=480 ± 7.5 nm, I=1.97 mW/cm^2^, 40 s, 4000 fl) and found that Opto-β_2_AR-All-In elicited a pronounced increase of intracellular cAMP levels (>50-fold above baseline) (**Fig. 2g,h**). This result is comparable to ligand-induced β_2_AR responses in this cell type (see below) and demonstrates that inclusion of structural contacts yielded a chimeric receptor that potently activates downstream signaling. We went on to further characterize the rationally-designed Opto-β_2_AR-All-In, which we termed Opto-β_2_AR-2.0, first in comparison to β_2_AR, then to a previous OptoXR and finally to variants that exhibit altered contacts.

**Fig. 2:**
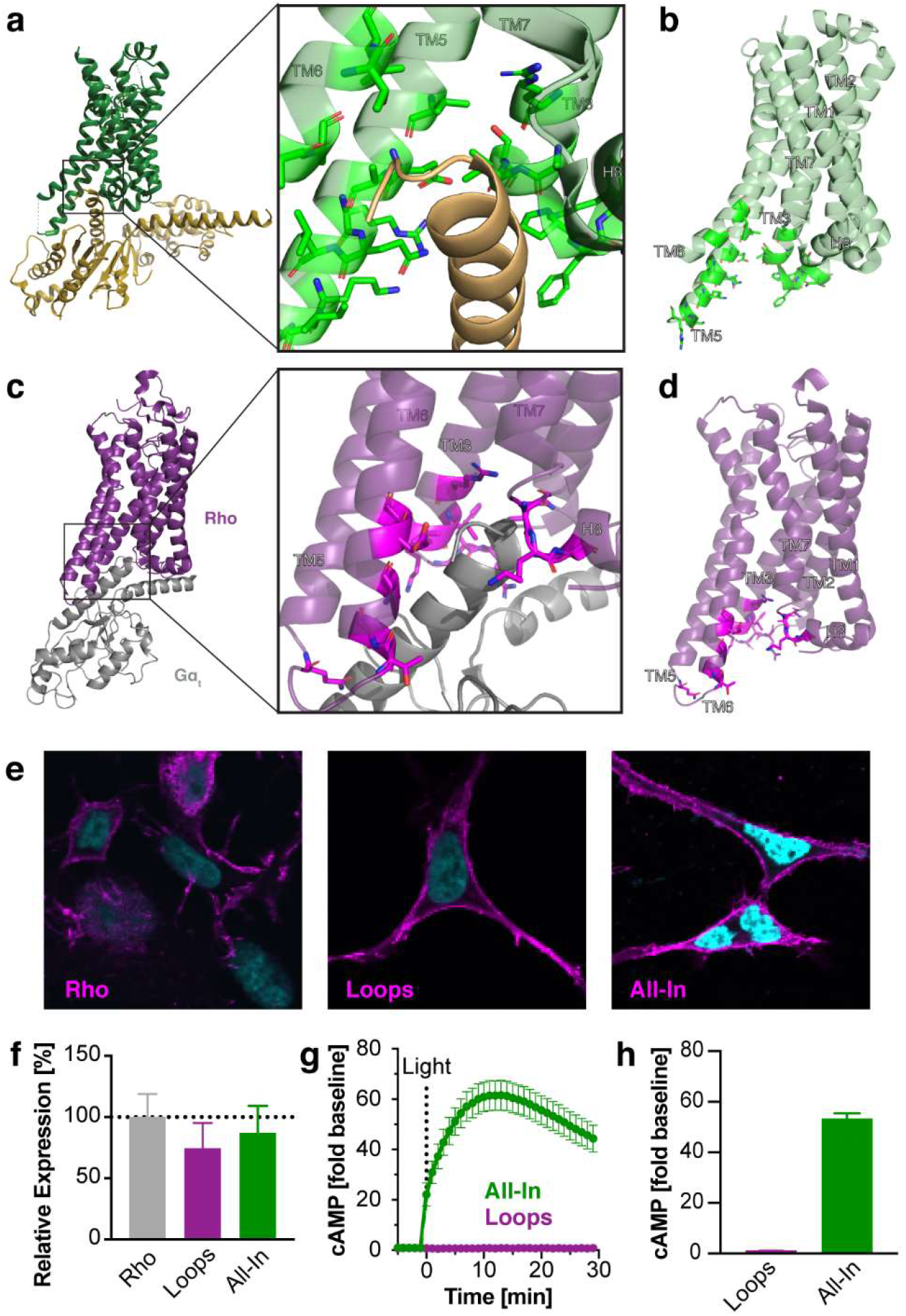
Identification of functionally-relevant domain boundaries. **(a-d)** Gα contacts on β_2_AR (a, b) and rho (c, d) defined as described in the main text (PDB ID: 3SN6 and 6OYA). Contacts extend upwards on TM α-helices. **(e)** Confocal microscopy of rho, Opto-β_2_AR-Loops and -All-In (N-terminal rho epitope 4D2). **(f)** FC analysis of rho, Opto-β_2_AR-Loops and -All-In expression using the same antibody as in (e). **(g)** Time course of cAMP production following light stimulation of OptoXRs (λ=480 ± 7.5 nm, I=1.97 mW/cm^2^, 40 s, 4000 fl). **(h)** Maximal cAMP signals elicited by OptoXRs (see (f) for light stimulation parameters). For f-h: n=9, 3 independent experiments. Data shown as mean ± S.E.M.

### Signaling profile of Opto-β_2_AR-2.0

We next characterized the signaling profile of Opto-β_2_AR-2.0 in comparison to β_2_AR for multiple signaling pathways (**Fig. 3a**). We first investigated intracellular cAMP production in more detail. HEK293 cells expressing Opto-β_2_AR-2.0 or β_2_AR were stimulated with light (λ=480 ± 7.5 nm, I=1.97 mW/cm^2^, 0.01-10 s, 1-4000 fl) or the potent β-adrenoreceptor agonist isoproterenol (ISO) followed by real-time cAMP detection (**Fig. 3b**). Generally, cAMP levels reached upon Opto-β_2_AR-2.0 activation were similar to those reached upon β_2_AR activation, and both receptors exhibited dose-dependent increases (half-maximal activation light dose (LD50) for Opto-β_2_AR-2.0: 17 fl (0.17 s, λ=480 ± 7.5 nm, I=1.97 mW/cm^2^, EC50 for β_2_AR: 8.7 nM ISO) (**Fig. 3c,d**). Stimulation of Opto-β_2_AR-2.0 and β_2_AR induced similar cAMP time courses (**Fig. 3b, Supplementary Fig. S4**).

**Fig. 3:**
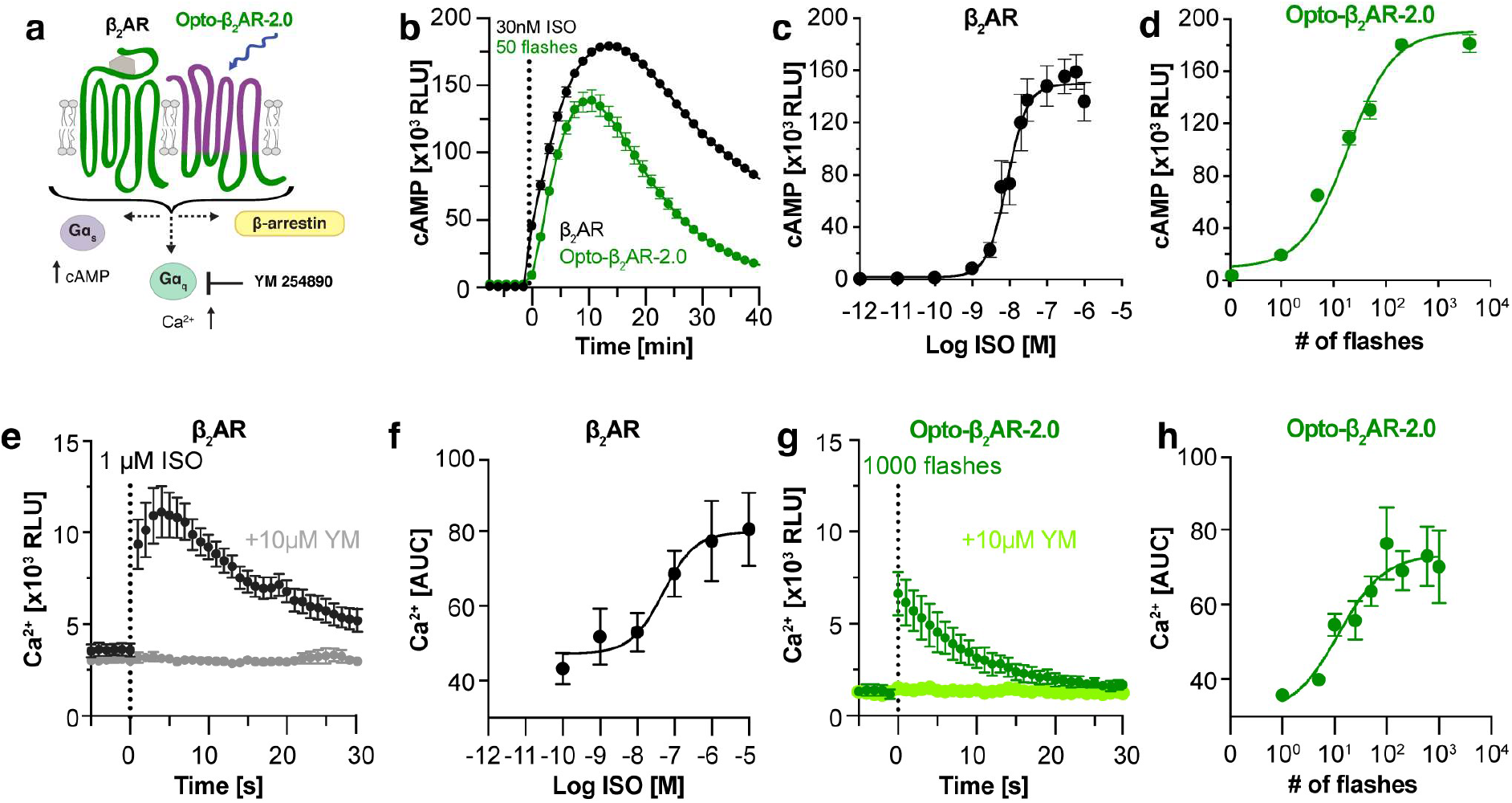
Signaling by Opto-β_2_AR-2.0. **(a)** Signaling pathways examined for β_2_AR and Opto-β_2_AR-2.0. **(b)** Time course of cAMP production following stimulation of β_2_AR with ISO (1 μM) or Opto-β_2_AR-2.0 with light (λ=480 ± 7.5 nm, I=1.97 mW/cm^2^, 0.5 s, 50 fl). **(c, d)** Dose-dependent cAMP production following stimulation of β_2_AR with ISO or Opto-β_2_AR-2.0 with light (λ=480 ± 7.5 nm, I=1.97 mW/cm^2^, 0.01-20 s, 1-2000 fl). **(e, g)** Time course of intracellular Ca^2+^ mobilization following stimulation of β_2_AR with ISO (as in b) or Opto-β_2_AR-2.0 with light (as in b). Signaling was abolished by the Gα_q_ inhibitor YM. **(f, h)** Dose-dependent Ca^2+^ mobilization following stimulation of β_2_AR with ISO or Opto-β_2_AR-2.0 with light (λ=480 ± 7.5 nm, I=1.97 mW/cm^2^, 0.1-10 s, 1-1000 fl). For c, d, f, h: n=9-12, 3-4 independent experiments. Data shown as mean ± S.E.M.

We also investigated the ability of Opto-β_2_AR-2.0 and β_2_AR to mobilize intracellular Ca^2+^ in response to light or ISO. Analogous to cAMP production, both receptors induced dose-dependent Ca^2+^ signals to similar maximal levels (**Fig. 3e-h**). As for cAMP production, reversal to basal Ca^2+^ levels was marginally more rapid for Opto-β_2_AR-2.0 than β_2_AR (**Fig. 3e,g**). GPCRs can induce intracellular Ca^2+^ mobilization through multiple pathways, including the canonical Gα_q_ axis. To test whether β_2_AR and Opto-β_2_AR-2.0 activate Ca^2+^ *via* this axis, we applied the Gα_q_ inhibitor YM254890 (YM)^57^ and found that it indeed abolished signaling downstream of both receptors (**Fig. 3e,g**).

The general agreement between Opto-β_2_AR-2.0 and β_2_AR signaling observed for G-protein dependent pathways did strikingly not extend to β-arrestin. β_2_AR, like many other GPCRs, exhibits the ability to induce outcomes downstream of G-proteins and β-arrestin in a ligand-dependent manner. We quantified β-arrestin2 recruitment to β_2_AR and Opto-β_2_AR-2.0 using sensitive bioluminescence resonance energy transfer assays (BRET) (**Supplementary Fig. S5a**). β_2_AR recruited β-arrestin2 in a dose-dependent manner with a half-maximal effective ISO dose similar to that observed for cAMP induction (EC50: 18 nM; **Supplementary Fig. S5b,c**). However, we did not observe any light-dependent β-arrestin recruitment for Opto-β_2_AR-2.0 even at high light doses (λ=480 ± 7.5 nm, I=1.97 mW/cm^2^, 40 s, 4000 fl) (**Supplementary Fig. S5d**) pointing to a preference of Opto-β_2_AR-2.0 for Gα-coupled pathways. It is noteworthy that β-arrestin recruitment has not been examined in earlier studies of β_2_AR OptoXRs (as discussed below). The result that β_2_AR and a β_2_AR OptoXR receptor exhibit distinct behavior may not be entirely surprising. TM domains contribute to active state conformations and kinetics and ultimately signaling bias^58–60^ and these domains are different in β_2_AR and the rho chimera. Collectively, we found striking similarities but also differences in in the signaling responses elicited by Opto-β_2_AR-2.0 and β_2_AR.

### Optical activation profile and comparison to earlier Opto-β_2_AR

We went on to characterize the optical activation profile of Opto-β_2_AR and also compared it to an earlier β_2_AR OptoXR^18,19,26^. We found that whilst light stimulation for 2 s (λ=480 ± 7.5 nm, I=1.97 mW/cm^2^, 200 fl) was required to produce saturating cAMP levels, even a single 10 ms light flash induced a signal that was sizeable (~10% of maximal cAMP induction) and transient (decay in 7.5 min and thus faster than the decay after application of the amplitude-equivalent dose of 10 nM ISO) (**Fig. 4a**, **Supplementary Fig. S4**). Opto-β_2_AR-2.0 employs rho as light-sensing photoreceptor core which is sensitive to blue-green light across a relatively broad wavelength range (maximal absorption wavelength: 500 nm, FWHM: ~100 nm)^61,62^. We confirmed the spectral sensitivity of Opto-β_2_AR-2.0 by stimulating cells with light of different wavelengths (**Fig. 4b**). As expected given the broad absorption peak, Opto-β_2_AR-2.0 could be activated at 480 ± 7.5 nm (the closest wavelength to 500 nm available to us; I=1.97 mW/cm^2^) as well as at 445 ± 7.5 nm and 540 ± 7.5 nm (I=1.75 and 1.35 mW/cm^2^, 0.01-10 s, 1-1000 fl).

**Fig. 4:**
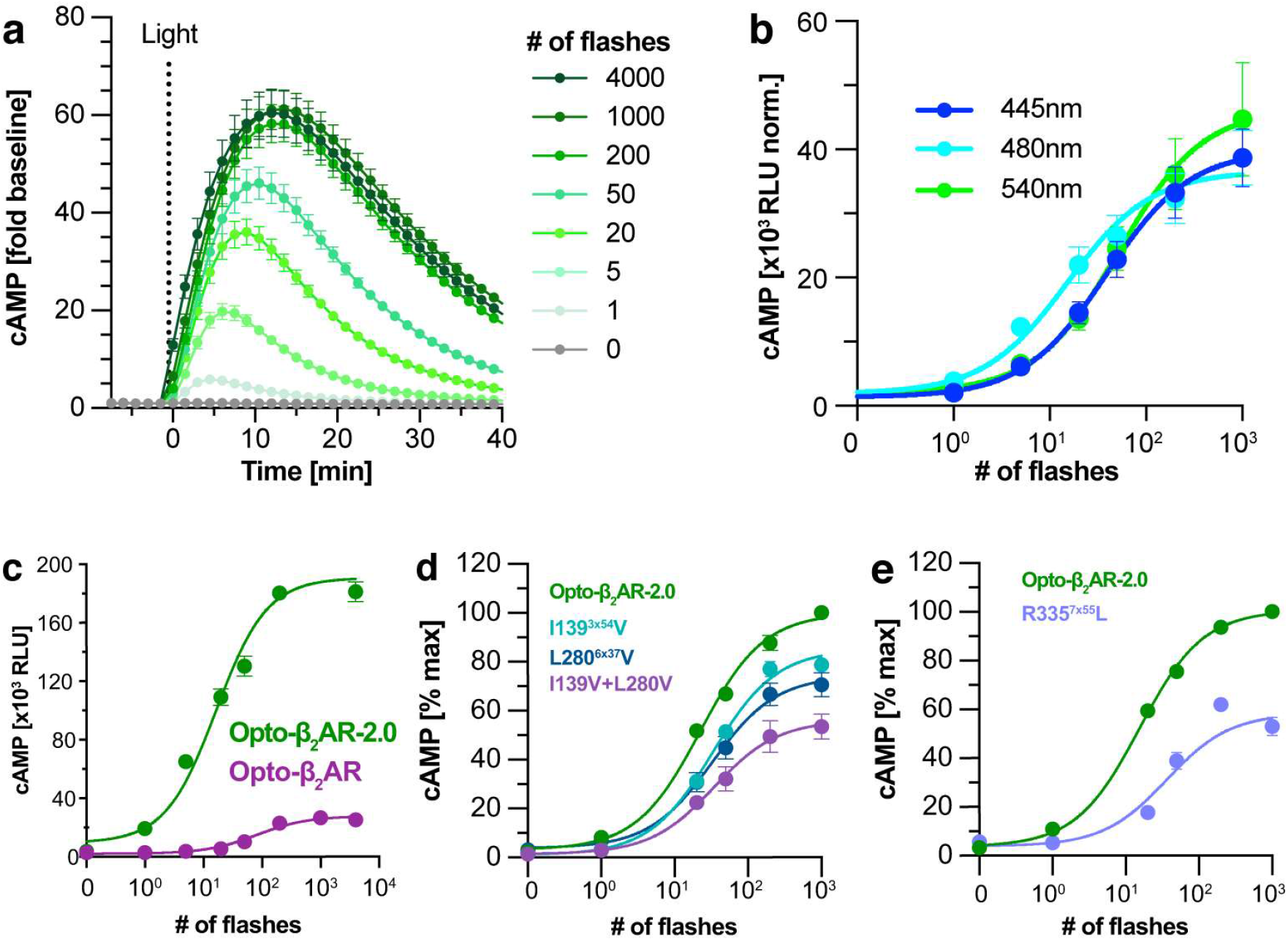
Optical activation of Opto-β_2_AR-2.0, Opto-β_2_AR and variant receptors. **(a)** Time course of cAMP production following stimulation of Opto-β_2_AR-2.0 (λ=480 ± 7.5 nm, I=1.97 mW/cm^2^, 0.01-40 s, 1-4000 fl). **(b)** Stimulation of Opto-β_2_AR-2.0 with light of different wavelengths (λ=480 ± 7.5 nm, I=1.97 mW/cm^2^; λ=445 ± 7.5 nm, I=1.75 mW/cm^2^; λ=540 ± 7.5 nm, I=1.35 mW/cm^2^; for all wavelengths: 0.01-10 s, 1-1000 fl). **(c)** Stimulation of Opto-β_2_AR and Opto-β_2_AR-2.0 with increasing light doses (λ=480 ± 7.5 nm, I=1.97 mW/cm^2^, 0.01-40 s, 1-4000 fl). **(d, e)** Stimulation of Opto-β_2_AR-2.0 reversal variants I139^3×54^V, L270^6×37^V and the corresponding double mutant (d) as well as R335^7×55^ (e) with light (λ=480 ± 7.5 nm, I=1.97 mW/cm^2^, 0.01-10 s, 1-1000 fl). For b-e: n=6-9, 2-3 independent experiments. Data shown as mean ± S.E.M.

In the seminal study of Kim *et al*.^19^, a β_2_AR OptoXR was designed using a screening approach and this receptor was later expressed *in vivo* and ex vivo^18,26^. Following the current convention, we refer to this earlier receptor as Opto-β_2_AR whilst our receptors carry further labels (e.g., the label 2.0; the domain boundaries of Opto-β_2_AR and Opto-β_2_AR-2.0 shown side-by-side in **Supplementary Fig. S3**). We compared the efficiency of Opto-β_2_AR-2.0 to Opto-β_2_AR and found that Opto-β_2_AR-2.0 generated larger maximal signals (by 7.2 fold) and produced half-maximal signals at 4.8-fold lower light doses (**Fig. 4c**) (LD50 of Opto-β_2_AR: 82 fl, 0.82s; LD50 of Opto-β_2_AR-2.0: 17 fl, 0.17s; in both cases λ=480 ± 7.5 nm, I=1.97 mW/cm^2^; expression levels were comparable, **Supplementary Fig. S6**). On the time scale of our longest repetition experiments (~1.5 h), cAMP signals generated by Opto-β_2_AR-2.0 were even comparable to the potent jellyfish opsin JellyOp^16^ (**Supplementary Fig. S7**). One further notable observation was that the activity of Opto-β_2_AR, but not that of Opto-β_2_AR-2.0 or β_2_AR, was sensitive to the Gα_i_ inhibitor PTX (**Supplementary Fig. S8**). This result suggests that the original Opto-β_2_AR exhibited a signaling activity that is in this cell type not representative of β_2_AR but may be relevant in the context of previous and future applications of this earlier receptor. Understanding the mechanism underlying this activity requires separate further study as it cannot be attributed to the absence/presence of structural GPCR-Gα contacts: (i) no rho-Gα_i/o/t_ contact residues remain in Opto-β_2_AR (**Supplementary Fig. S3**) and (ii) Gα_s_ and Gα_i_ associate with the homologous β_1_AR receptor through an overlapping ensemble of residues^63^ all of which are present in β_2_AR and Opto-β_2_AR-2.0. While the effect of PTX was pronounced, cAMP production by Opto-β_2_AR even under conditions of Gα_i_ inhibition was still several fold below that of Opto-β_2_AR-2.0 (**Supplementary Fig. S8**). Collectively, these data show that Opto-β_2_AR-2.0 is an efficient optical actuator that offers higher potency and specificity than its closest known OptoXR relative.

### Contributions of contact residues to Opto-β_2_AR-2.0 function

We next tested if the functional contribution of individual structural contacts can be quantified in Opto-β_2_AR-2.0. We examined key residues that (i) are located in the three main receptors domains that were exchanged between rho and β_2_AR (TM3-ICL2-TM4, TM5-ICL3-TM6 and TM7-C), (ii) form contacts (e.g., hydrophobic interactions or charge-charge interactions) to Gα_s_ and (iii) are present in Opto-β_2_AR-2.0 but absent in Opto-β_2_AR. In the domains TM3-ICL2-TM4 and TM5-ICL3-TM6, we focused on I139^3×54^ and L270^6×37^ that form hydrophobic contacts to Gα_s_ (in both cases through Cδ side chain atoms^36^; residue numbers correspond to the Opto-β_2_AR-2.0 sequence and, in superscript, to the GPCRdb numbering scheme^64^). Upon substitution of these residues to the corresponding rho residues present in Opto-β_2_AR (I139^3×54^V and L270^6×37^V; Val does not contain a Cδ atom), we found reduced cAMP induction by 15 and 25% as well as ~2-fold reduced LD50 values (33.3 fl, 0.34 s and 32.9 fl, 0.33 s, respectively; in both cases λ=480 ± 7.5 nm, I=1.97 mW/cm^2^) (**Fig. 4d**). A I139^3×54^V L270^6×37^V double mutant resulted in further loss of function (LD50 34.4 flashes, 0.35 s, 44% reduced cAMP induction) (**Fig. 4d**). We also investigated TM7. Specifically, residue R335^7×55^ at the cytoplasmic end of this TM α-helix has been implicated in Gα_s_-binding through a charge-charge contact to residue E392^H5.24^ of Gα_s_ in structural studies and molecular dynamics simulations^43,65^. A deleterious substitution (R335L^7×55^; modelled after a naturally observed receptor variant^66^) resulted in maximal cAMP levels that were reduced by 42% (LD50: 36.6 fl, 0.37s, λ=480 ± 7.5 nm, I=1.97 mW/cm^2^) (**Fig. 4e**). Overall, the function of Opto-β_2_AR-2.0 is indeed sensitive to substitution of newly-introduced residues that were identified in structural GPCR-Gα complex analysis.

### Structure-guided receptors as a development platform

We went on to demonstrate consequences of the optimization approach. Above exploration of contact residues was assisted by the large signals generated by Opto-β_2_AR-2.0: A quantitative analysis could still be performed despite a marked reduction in activity because of residue substitutions. We reasoned that the potency of Opto-β_2_AR-2.0 may also enable experiments where the photosensitive domain of the receptor is engineered as also this manipulation may be associated with reduced function. Of importance in optogenetic tool development is an availability of actuator proteins with sensitivities to different wavelengths (“blue-shifting” (towards lower maximal absorption wavelengths) or “red-shifting” (towards higher maximal absorption wavelengths))^67–70^. This is generally achieved either through identification of naturally-occurring spectrally-diverse photoreceptors, e.g. cone opsins in the case of GPCRs^11^, or through tuning of photoreceptors using substitutions in their chromophore binding pockets^71,72^. Binding pocket mutagenesis can be associated with reduced function in opsins^73–76^, which is one possible reason why spectral tuning using mutagenesis has never been demonstrated in OptoXRs. Indeed, when introducing three substitutions in the original Opto-β_2_AR that were previously shown to blue-shift rho absorption by ~50 nm^75^ (T118^3×33^A, E122^3×37^D and A318^7×39^S; residue numbers correspond to Opto-β_2_AR-2.0/GPCRdb numbering; **Fig. 5a**), no light-induced activity could be detected (λ=480 ± 7.5 nm, I=1.97 mW/cm^2^, 0.01-10 s, 1-1000 fl) (**Fig. 5b**). In turn, introduction of the same residues in Opto-β_2_AR-2.0 resulted in a blue-shifted variant (termed Opto-β_2_AR-2.0-Blue) that induced cAMP to levels that were comparable to previous non-shifted OptoXRs (λ=480 ± 7.5 nm, I=1.97 mW/cm^2^, 0.01-10 s, 1-1000 fl) (**Fig. 5c**). As expected, in the spectrally tuned Opto-β_2_AR-2.0-Blue sensitivity to blue light was maximal and to green light reduced (**Fig. 5c**).

**Fig. 5:**
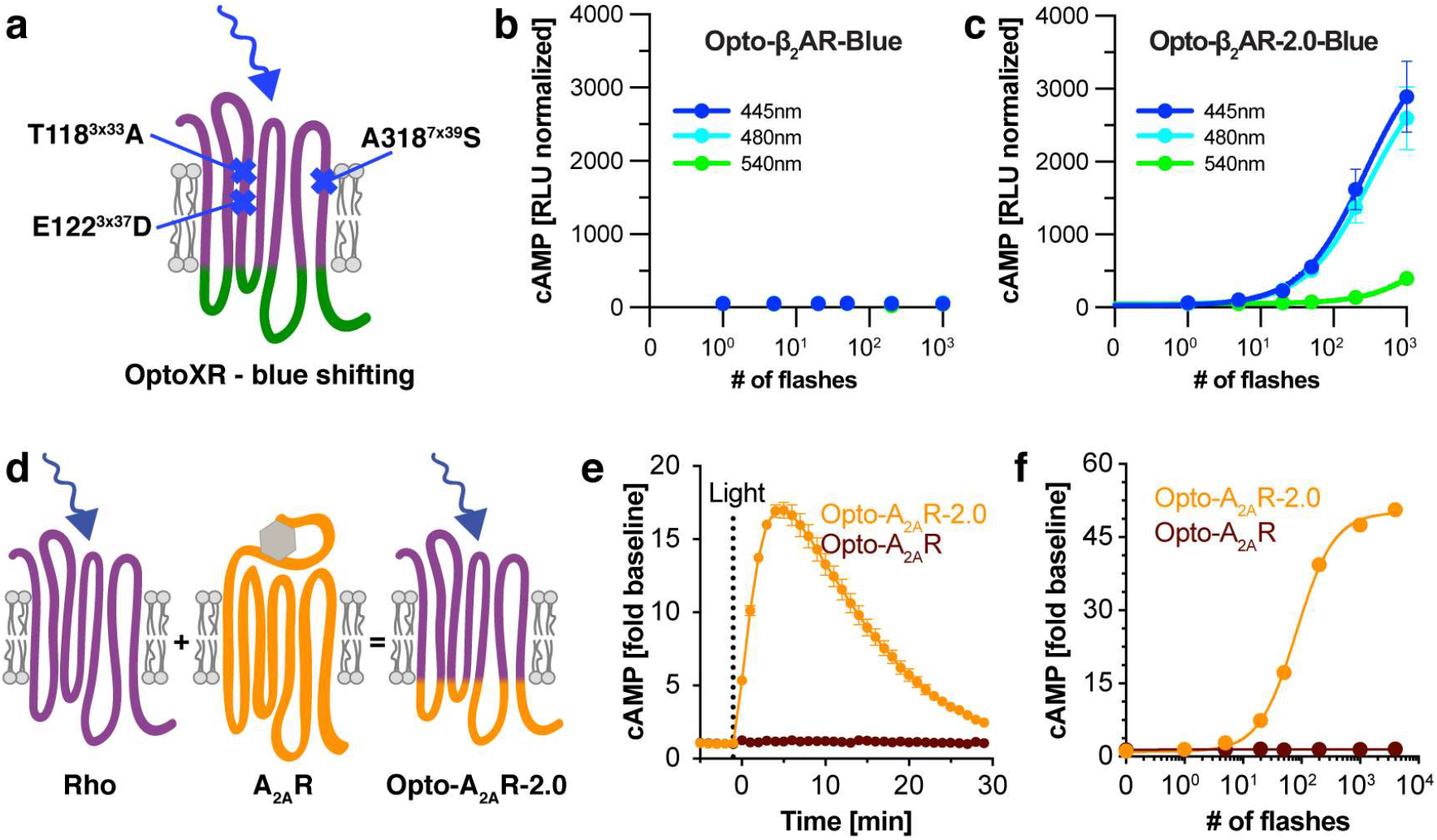
Diversified receptor variants. **(a)** Substitutions in Opto-β_2_AR and Opto-β_2_AR-2.0 to engineer variants termed Opto-β_2_AR-Blue and Opto-β_2_AR-2.0-Blue. **(b,c)** Stimulation of Opto-β_2_AR-Blue and Opto-β_2_AR-2.0-Blue with light of different wavelengths (λ=480 ± 7.5 nm, I=1.97 mW/cm^2^; λ=445 ± 7.5 nm, I=1.75 mW/cm^2^; λ=540 ± 7.5 nm, I=1.35 mW/cm^2^; for all wavelengths: 0.01-10 s, 1-1000 fl). **(d)** Opto-A_2A_R-2.0. **(e)** Time course of cAMP production following stimulation of Opto-A_2A_R and Opto-A_2A_R-2.0 with light (λ=480 ± 7.5 nm, I=1.97 mW/cm^2^, 0.5 s, 50 fl). **(f)** Dose-dependent cAMP production following stimulation Opto-A_2A_R-2.0 and Opto-A_2A_R with light (λ=480 ± 7.5 nm, I=1.97 mW/cm^2^, 0.1-20 s, 1-2000 fl). For panels b, c, f: n=6-9, 2-3 independent experiments. Data shown as mean ± S.E.M.

We also explored whether a GPCR-Gα contact-based redesign may be applicable to a further receptor (**Fig. 5d**). We have previously developed a light-activated A_2A_R (Opto-A_2A_R) and characterized its function in highly-sensitive because integrative transcriptional reporter assays^47^. Analogously as for Opto-β_2_AR-2.0 above, we determined G-protein proximal residues in the A_2A_R receptor structure^38^ to inform the placement of domain boundaries following the “All-In”-rationale. We termed the resulting receptor Opto-A_2A_R-2.0. Opto-A_2A_R-2.0 also contained additional A_2A_R residues in TM3, TM6 and the C-terminus when compared to Opto-A_2A_R (**Supplementary Tables S3 and S4**). We went on to test the ability of both receptors to induce cAMP responses and found that only Opto-A_2A_R-2.0 resulted in detectable activation in the real-time assays (**Fig. 5e,f**). Thus the structure-guided approach permitted generation of a functionally improved A_2A_R OptoXR.

## DISCUSSION

GPCRs relay diverse extracellular signals to specific downstream signaling pathways and thereby regulate essential physiology. To control GPCR signaling with spatio-temporal precision, chimeric light-activated OptoXRs have been developed and deployed alongside repurposed naturally-occurring opsins^20,21^. The design of most published OptoXRs relied on domain boundaries that were proposed in a seminal study^19^. Notably, this recipe was formulated at a time when few high-resolution structures were available for GPCRs and none for GPCR-G-protein complexes. The past fifteen years have seen the emergence of detailed structures depicting >150 unique GPCRs and >50 unique GPCR-signaling-protein complexes^1,30^. Here, we utilized some of this structural information for rational improvement of OptoXRs. First, we evaluated if sequence alignments and structure alignments yield the same chimeric receptor backbones. For our set of prototypical receptors and domain boundaries this was the case. Second, we identified receptor residues that are proximal to Gα-subunits to position the domain boundaries in modified chimeric receptors. We found that an “All-In”-rationale, which incorporates the three ILCs, the C-terminus and the identified structural GPCR-Gα contact regions, markedly improved the function of a β_2_AR OptoXR. These modifications, along with a non-functional chimeric receptor that contained only the ICLs of β_2_AR, demonstrated how TM α-helical residues contribute to efficient downstream coupling. As we incorporated all residues identified in the complex structure, a rationale for further optimization is not directly evident (see below for a test of the contribution of inserted residues). The approach we have taken is structure-based and thus fundamentally different from, e.g., the screening-based methods employed on OptoXRs previously or on GPCR-based fluorescent sensors. In Opto-β_2_AR-2.0, many of the incorporated TM α-helical residues (e.g., I139^3×54^, L280^6×37^ or R335^7×55^) are conserved in Gα_s_-coupled aminergic (including adrenergic) receptors but not in non-Gα_s_-coupled receptors. Our comparison of Opto-β_2_AR-2.0 and a previous Opto-β_2_AR, which retained rho residues at the corresponding sites, and a reverse experiment in which we substituted these residues in Opto-β_2_AR-2.0, confirmed their contributions to function. In addition, our analysis also rationalizes previous findings that omission of larger structural domains of β_2_AR, e.g. entire intracellular domains, resulted in reduced activity of chimeric GPCRs^16,19,27^.

Our work is complementary to two elegant recent studies in which coupling downstream of GPCRs was altered using structural information. Using simulations of terminal Gα α-helices in combination with functional assays, Sandhu et al. generated a triple mutant adrenoreceptor with more promiscuous downstream coupling, in particular leading to more potent activation of Gα_q_ compared to the WT receptor^43^. Using structure modeling and *in silico* and experimental mutagenesis, Young et al. engineered orthogonal receptor-Gα-pairs that interacted specifically and independently of their endogenous counterparts^42^. Collectively, these studies and our work demonstrate separate advances towards several design objectives (coupling switch, promiscuity or orthogonality) using distinct strategies (chimeric receptors or mutagenesis).

One element of our study was examination to which degree this OptoXR recapitulates the function of the ligand-activated target receptor. GPCRs assume distinct ligand-dependent active states and state lifetimes that lead to preferred activation of some of the available signaling pathways over others^58–60,77–79^. OptoXR activation likewise was associated with a particular signaling profile. Given pronounced differences between rho and β_2_AR, it may not be surprising that the signaling profile of Opto-β_2_AR-2.0 was not identical to that of β_2_AR. In particular, we were unable to detect β-arrestin recruitment for Opto-β_2_AR-2.0 and thus under the conditions examined here the activity of Opto-β_2_AR-2.0 resembled that of β_2_AR in response to G-protein-biased ligands, such as pepducins^80^. It is noteworthy that an ability to predominantly activate G-protein pathways may be desirable in some physiological scenarios^80,81^. This result may also be relevant in the context of an earlier study that proposed the development of a “functionally-selective” Opto-β_2_AR with limited Gα_s_ signaling but induction of pathways that are known to be downstream of β-arrestin^26^. Notably, β-arrestin recruitment was not explored in this study and selectivity was achieved through mutagenesis of key functional motifs which were (i) located within the rho core and (ii) present in most Class A GPCRs (e.g., the highly-conserved Tyr^3×51^ of the E/DRY motif or a further tyrosine (Tyr^5×58^) that participates in active-state-specific interactions)^26,31^. Because of the placement of these substitutions in rho domains (as opposed to in β_2_AR-derived domains as here), a comparison of the previous results and ours is not possible.

The large signaling response elicited by Opto-β_2_AR-2.0 prompted us to explore further capabilities of OptoXRs. We introduced a spectral absorption shift to the rho core of Opto-β_2_AR-2.0 to diversify function. The blue-shifted receptor may complement blue-light sensitive cone opsins that activate opposing signaling responses (Gα_i/o_ instead of Gα_s_)^11^. Mutagenesis in binding pockets are associated with reduced protein folding and function in many protein families including opsins^73–76^. Indeed, we observed that substitution of three residues in proximity of the retinal cofactor impacted the function of the original Opto-β_2_AR such that we were not able to detect changes in cAMP levels. In contrast, Opto-β_2_AR-2.0-Blue induced changes that were even comparable to those produced by earlier unperturbed OptoXRs. Finally, again prompted by the functional improvement observed in Opto-β_2_AR-2.0, we went on to show that a structure-guided approach can also be applied to generate Opto-A_2A_R-2.0.

Structural biology has had a profound impact on the field of optogenetics, e.g. by enabling the rational design of light-activated ion channels, enzymes and protein-protein interactions^71,73,82^. We show that rational exchange of receptor domains resulted in more potent OptoXRs and enabled OptoXRs with new functions. However, because this is the first demonstration of structure-guided coupling engineering in OptoXRs, and one of very few in GPCRs in general^42,43^, further studies are required to test how universal this approach is. It is interesting to note that the temporal sequence of events driving the association of the G-protein to a GPCR is an active area of study. It is possible that this association initially involves intermediate receptor states and consequently contacts other than those observed in the stable GPCR-G-protein complexes; however, such contacts have not yet been identified^65,83,84^ and thus cannot be incorporated in a receptor design. OptoXRs may in the future support time-resolved studies of these binding events by providing non-invasive activation tools, similarly to successful photoreceptor applications in spectroscopy or drug discovery^85,86^. In addition to serving as an experimental platform, targeted OptoXRs may also continue to make contributions towards understanding the impact of signaling on cells and cellular networks.

## AUTHOR CONTRIBUTIONS

Conceptualization, A.M.T., E.J.G. and H.J.; Funding Acquisition, E.J.G. and H.J.; Methodology, A.M.T., W.L.S., E.J.G. and H.J.; Project Administration, H.J.; Investigation, A.M.T., W.L.S. and E.J.G.; Data curation, A.M.T., W.L.S., E.J.G. and H.J.; Supervision, E.J.G. and H.J.; Writing - Original Draft, A.M.T. and H.J.; Writing - Review and Editing, H.J.

## ACKNOWLEDGEMENTS

We thank M. Halls and P. Sexton for materials and advice. This study was supported by grants of the Australian Research Council (DP200102093, to H.J.), the National Health and Medical Research Council (APP1187638, to H.J.) and a fellowship of the Synthetic Biology Future Science Platform of the Commonwealth Scientific and Industrial Research Organisation (to E.J.G.). The Australian Regenerative Medicine Institute is supported by grants from the State Government of Victoria and the Australian Government. The EMBL Australia Partnership Laboratory (EMBL Australia) is supported by the National Collaborative Research Infrastructure Strategy (NCRIS) of the Australian Government. MicroMon of Monash University provided Sanger sequencing services.

## CONFLICT OF INTEREST

The authors declare no conflict of interest.

## Notes

### Competing Interest Statement

The authors have declared no competing interest.

