## Supplemental Materials for "Structure-guided optimization of light-activated chimeric G-protein coupled receptors"

### SUPPLEMENTARY INFORMATION

Tichy *et al.*

**Contents:**

Supplementary Figures S1 to S8

Supplementary Tables S1 to S4

Supplementary References

#### SUPPLEMENTARY FIGURES

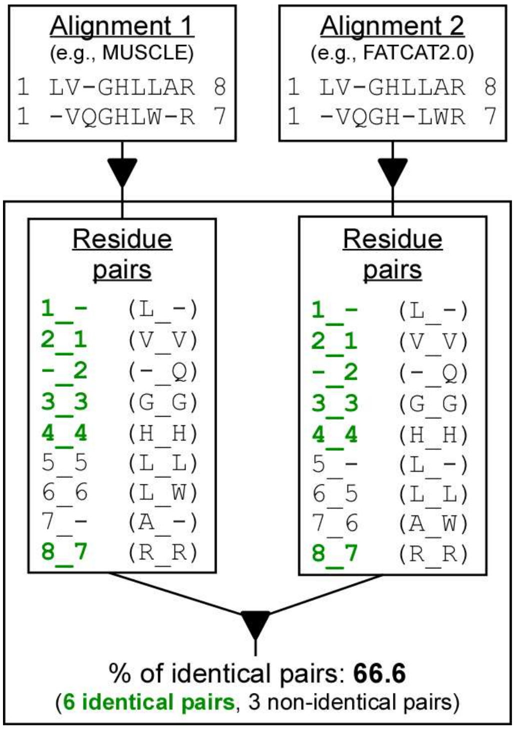


**Fig. S1: Procedure for quantitative comparison of two alignments.** Alignments are generated individually (e.g., using MUSCLE and FATCAT2.0) followed by comparison using this method (here, exemplary sequences are shown). For each alignment and each alignment position in the alignment, a residue pair in the format A_B is determined (where A is the number of the residue in the first sequence that aligns with residue B in the second sequence; e.g., 8_7 is the final residue pair of both exemplary alignments). The residue pairs are deposited in lists that are then compared to derive the percentage of identical pairs independent of their position in the list. Here, identical pairs are shown in green.

**a**

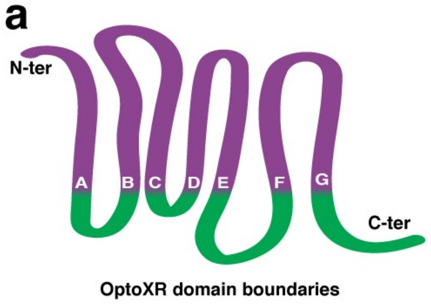


**b**

| | Difference between sequence- and structure-based alignments (# of amino acids) | | | |
|---|---|---|---|---|
| Domain boundary | Rho:$\beta_2$AR | Rho:$A_{2A}$R | Rho:$\beta_1$AR | Rho:$M_1$R |
| A | 0 | 0 | 0 | 0 |
| B | 0 | 0 | 0 | 0 |
| C | 0 | 0 | 0 | 0 |
| D | 0 | 0 | 0 | 0 |
| E | 0 | 0 | 1 | 0 |
| F | 0 | 0 | 1 | 0 |
| G | 0 | 0 | 0 | 0 |

**Fig. S2: Domain boundaries and comparison of sequence- and structure-based alignments at boundaries. (a)** Graphical representation of the seven domain boundaries (labelled A to F) in OptoXRs. **(b)** We tested if a sequence-based method (MUSCLE) and a structure-based method (FATCAT2.0) align identical residues at the seven boundaries. This comparison was performed for four prototypical receptor pairs and the prototypical boundaries proposed by Kim *et al.*[1]. Differences at the boundaries between these alignments are given in the table: A value of zero denotes that identical residues were aligned using the sequence- and structure-based method, whilst a value of one denotes a shift in alignment position by one residue. Generally, no differences were observed between these alignments methods with the exception of single residue deviations for Rho:$\beta_1$AR at boundary E and F.

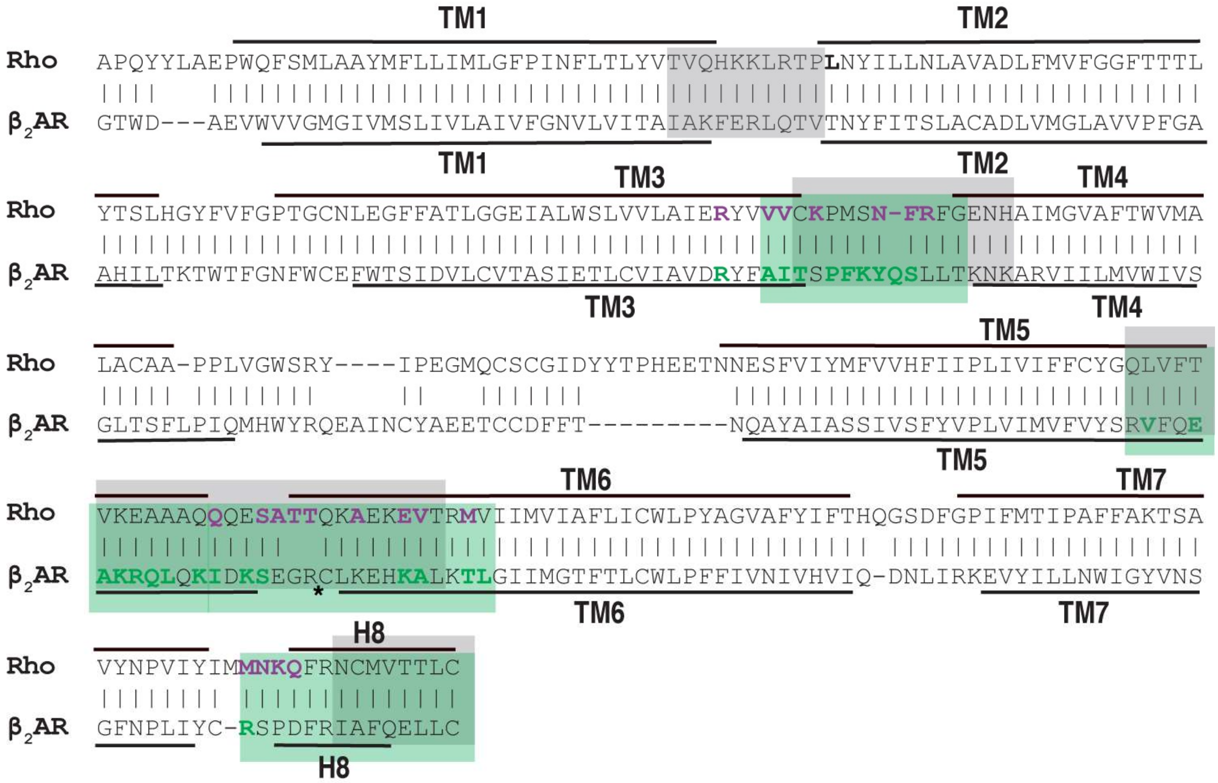


**Fig. S3: Structure-based alignment and contact residues.** Alignment of rho and $\beta_2$AR structures using FATCAT2.0 (PDB IDs: 6OYA (rho) and 3SN6 ($\beta_2$AR)). Pairs of residues that are structurally equivalent are connected by a vertical line. Bold residues: Structural contacts identified in multiple receptors (also see Supplementary Table S2 for a list of contacts). Grey highlights: Exchanged residues to generate Opto-$\beta_2$AR[1]. Green highlights: Exchanged residues to generate Opto-$\beta_2$AR-All-In/2.0. The star denotes the position of $\beta_2$AR ICL3 which is not resolved in structures. Not shown here are the unstructured receptor N- and C-termini (see Supplementary Tables S3 and S4 for complete receptor sequences).

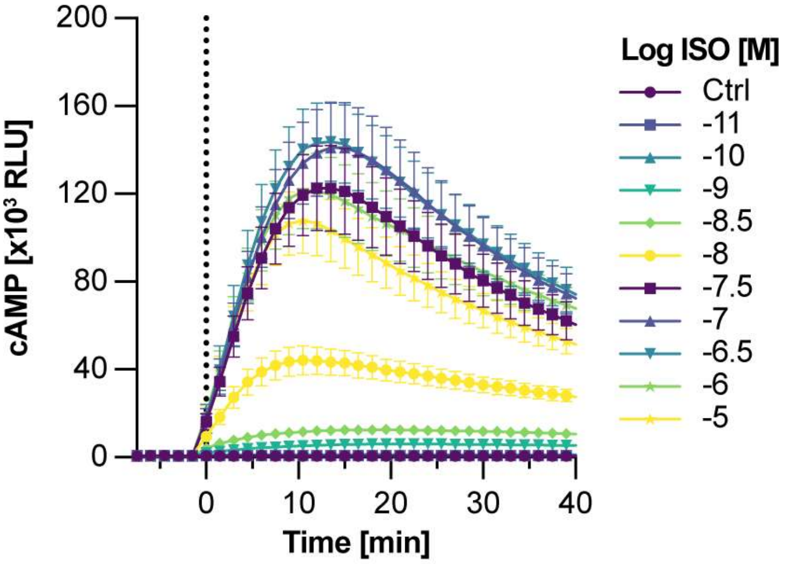


**Fig. S4: Time course of cAMP production after $\beta_2$AR stimulation.** n=9, 3 independent experiments. Data shown as mean ± S.E.M. Vertical line indicates light stimulation.

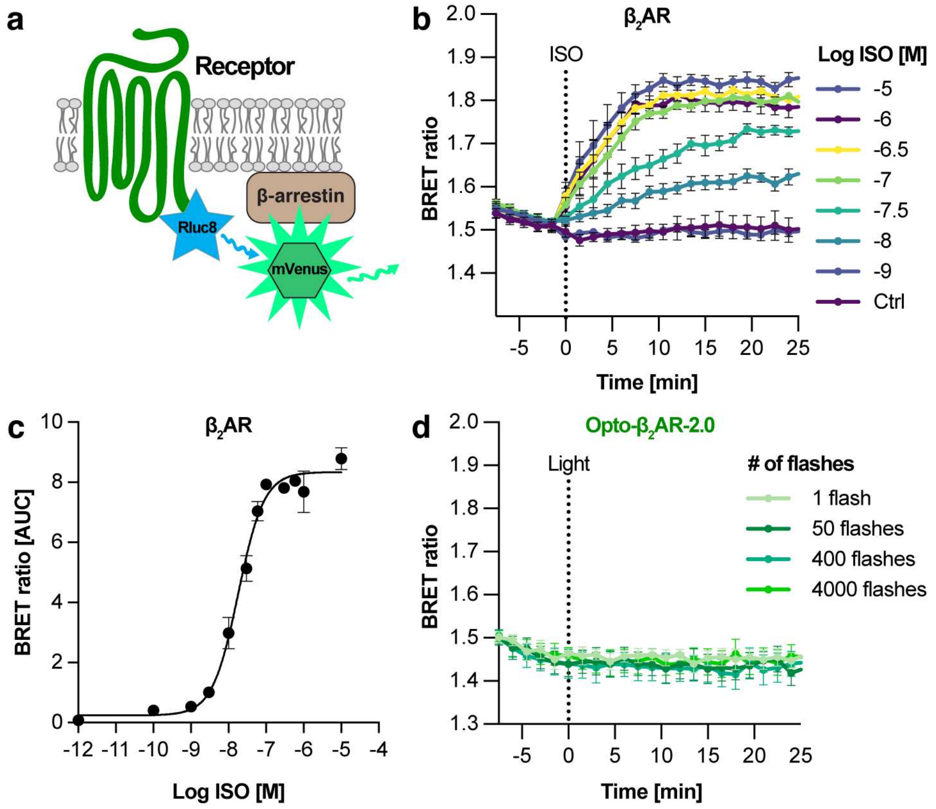


**Fig. S5: β-arrestin2 recruitment assay. (a)** Schematic of BRET assay to measure β-arrestin2 recruitment to Rluc8-tagged receptors. **(b)** Average time course of β-arrestin2 recruitment following stimulation of $\beta_2$AR with ISO. **(c)** Dose-dependent β-arrestin2 recruitment determined as area under the curve (AUC). **(d)** Average time course of β-arrestin2 recruitment following stimulation of Opto-$\beta_2$AR-2.0 with light (λ=480 ± 7.5 nm, I=1.97 mW/cm$^2$, 0.01-40 s, 1-4000 fl). For all panels: n=6-9, 2-3 independent experiments. Data shown as mean ± S.E.M.

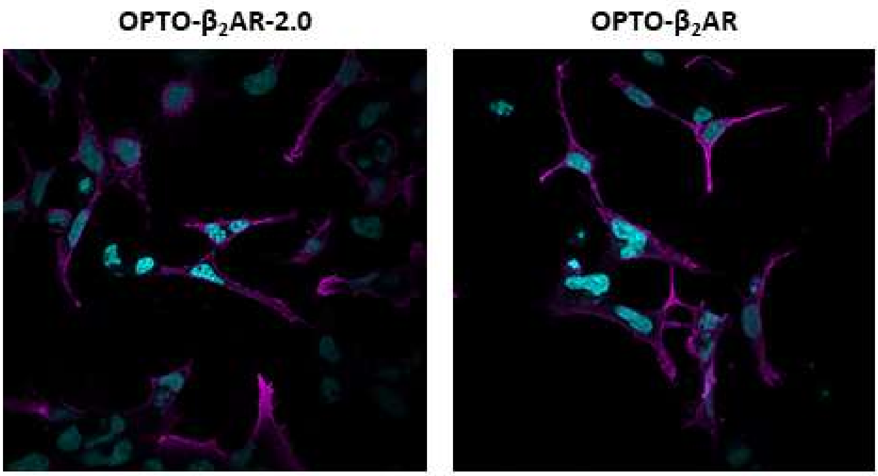


**Fig. S6: Confocal microscopy of Opto-β2AR-2.0 and Opto-β2AR.** Receptors were detected using the same antibody (N-terminal rho epitope 4D2).

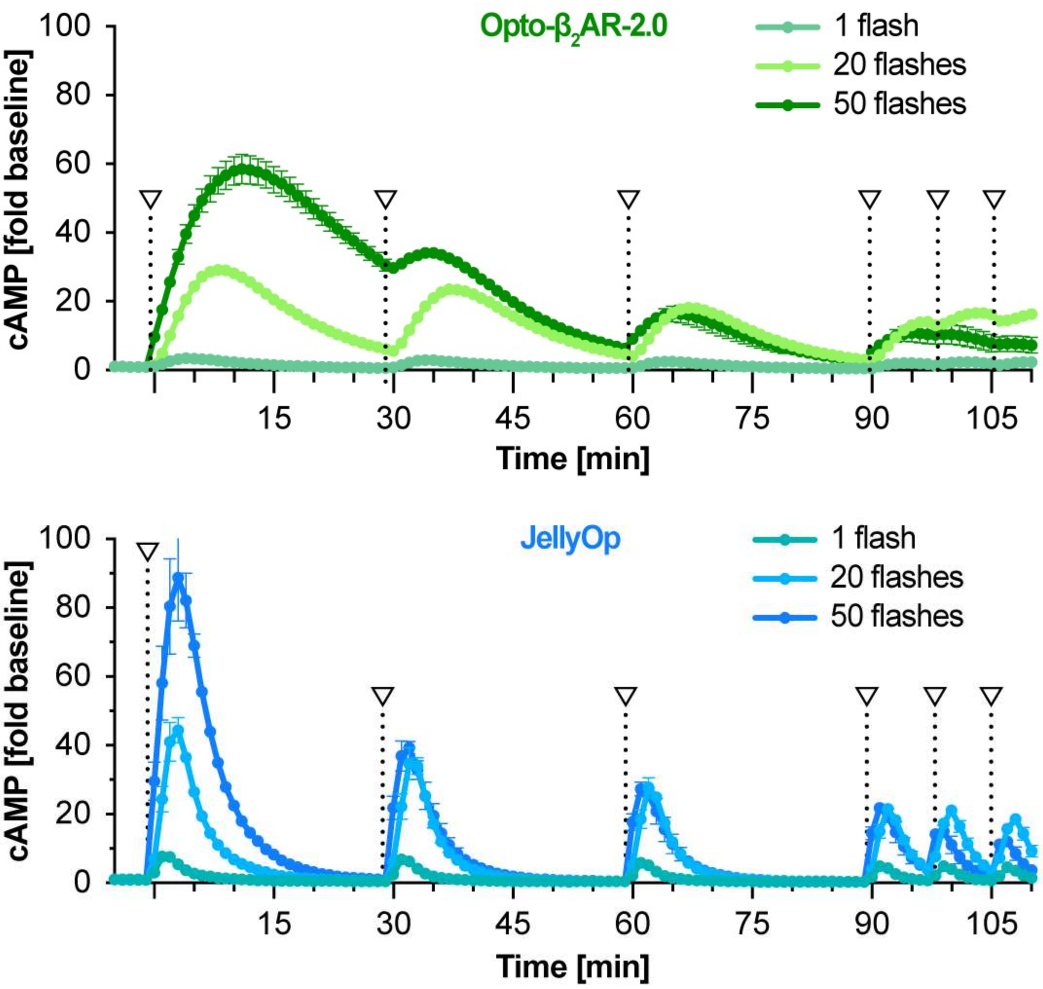


**Fig. S7: cAMP production following repeated light stimulation of Opto-$\beta_2$AR-2.0 (top) and JellyOp (bottom).** Light stimulation was at λ=480 ± 7.5 nm with I=1.97 mW/cm$^2$ for 0.01-0.5 s (1-50 fl). n=6, 2 independent experiments. Data shown as mean ± S.E.M.

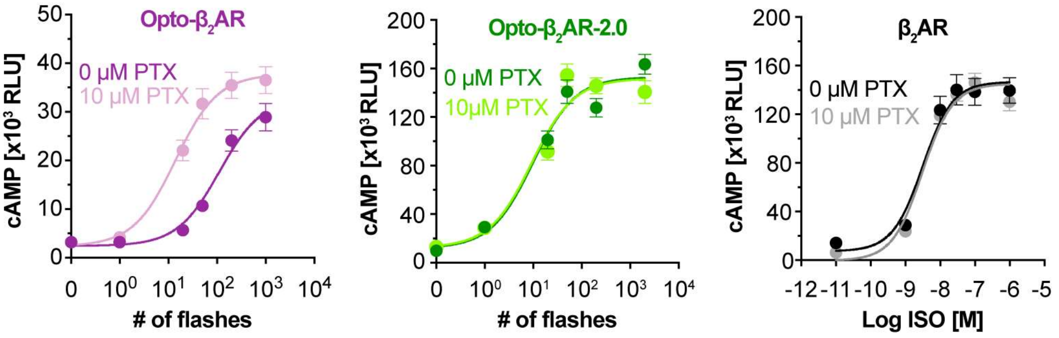


**Fig. S8: cAMP production following treatment with 10 µM PTX.** Opto-$\beta_2$AR and Opto-$\beta_2$AR-2.0 were stimulated with light ($\lambda$=480 ± 7.5 nm, I=1.97 mW/cm$^2$, 0.01-10 s, 1-1000 fl) and $\beta_2$AR with ISO. n=6-15, 2-5 independent experiments. Data shown as mean ± S.E.M.

#### SUPPLEMENTARY TABLES

**Table S1: A comprehensive list of previous OptoXRs with the respective methods for identification of domain boundaries and alignments.** For less common receptors, the following abbreviations were used: SW: short-wavelength opsin, JellyOP: opsin of the jellyfish *Carybdea rastonii*, OPN4: melanopsin, LW: long-wavelength opsin.

| Chimeric receptor | Domain boundaries identical to Kim *et al.*? | Alignment method | Reference |
|---|---|---|---|
| Rho:$\beta_2$AR | Reference study | Sequence alignment | 1 |
| Rho:$\beta_2$AR, Rho:$\alpha_1$AR | Yes | Sequence alignment | 2 |
| Rho:5-$HT_1$A | Yes | Sequence alignment | 3 |
| Rho:$\beta_2$AR | Yes | Sequence alignment | 4 |
| SW:JellyOp | Yes | Sequence alignment | 5 |
| Rho:µOR | Yes (with the exception of one residue in C-terminus) | Sequence alignment | 6 |
| Rho:D1R | Yes | Sequence alignment | 7 |
| Rho:CXCR4 | Yes | Sequence alignment | 8 |
| Rho:µOR | Yes (with the exception of one residue in ICL2 and two residues in ICL3) | Sequence alignment | 9 |
| Rho:$A_{2A}$R | Chimeric receptor sequence not available. | Unknown | 10 |
| OPN4:mGlu$R_6$ | No. The target receptor was a Class C GPCR. | Sequence alignment | 11 |
| OPN4:5-$HT_{2A}$ | No. Origin of boundaries unclear. | Sequence alignment | 12 |
| Rho:$\beta_2$AR, Rho:$\alpha_1$AR, Rho:$A_{2A}$R, Rho:$D_1$R, Rho:$D_2$R, Rho:$M_1$R, Rho:$M_2$R, Rho:$M_3$R, Rho:FFR3 | Yes | Sequence alignment | 13 |
| 63 orphan GPCRs | Yes | Sequence alignment | 13 |
| Rho:OPN4<br>LW:OPN4 | No. The authors focused on transferring ICLs and thus fewer residues than for boundaries of Kim *et al.* | Sequence alignment | 14 |

**Table S2: G-protein contacts determined for rho, β$_2$AR and A$_{2A}$R.** Grey: Receptor residues determined within 4 Å of Gα. Note: Residues of ICL3 are not listed here because they were not resolved in the respective structures (but these are included in the chimeric receptors). If available, multiple structures for each receptor were analyzed. PDB identifiers were for rho: 6CMO, 6OYA, for β$_2$AR: 3SN6, 7BZ2, and for A$_{2A}$R: 6GDF. Residue numbers follow the Class A GPCRdb numbering.

| GPCRdb$_A$ numbering | Rho (human) | β$_2$AR (human) | A$_{2A}$R (human) |
|---|---|---|---|
| TM2 | | | |
| 2x39 | L72 | T68 | T41 |
| TM3 | | | |
| 3x50 | R135 | R131 | R102 |
| 3x53 | V138 | A134 | A105 |
| 3x54 | V139 | I135 | I106 |
| 3x55 | C140 | T136 | R107 |
| 3x56 | K141 | S137 | I108 |
| ICL2 | | | |
| 34x50 | P142 | P138 | P109 |
| 34x51 | M143 | F139 | L110 |
| 34x52 | S144 | K140 | R111 |
| 34x53 | N145 | Y141 | Y112 |
| 34x54 | F146 | Q142 | N113 |
| 34x55 | R147 | S143 | G114 |
| TM5 | | | |
| 5x61 | L226 | V222 | I200 |
| 5x64 | T229 | E225 | A203 |
| 5x65 | V230 | A226 | A204 |
| 5x66 | K231 | K227 | R205 |
| 5x67 | E232 | R228 | R206 |
| 5x68 | A233 | Q229 | Q207 |

| | | | |
|---|---|---|---|
| 5x69 | A234 | L230 | L208 |
| 5x71 | Q236 | K232 | Q210 |
| 5x72 | Q237 | I233 | M211 |
| 5x74 | - | K235 | S213 |
| 5x75 | - | S236 | - |
| TM6 | | | |
| 6x23 | S240 | - | A221 |
| 6x24 | A241 | S262 | R222 |
| 6x25 | T242 | K263 | S223 |
| 6x26 | T243 | F264 | T224 |
| 6x29 | A246 | K267 | K227 |
| 6x32 | E249 | K270 | H230 |
| 6x33 | V250 | A271 | A231 |
| 6x36 | M253 | T274 | S234 |
| 6x37 | V254 | L275 | L235 |
| TM7 | | | |
| 7x55 | M308 | R328 | Y290 |
| 7x56 | M309 | - | R291 |
| H8/C-term | | | |
| 8x47 | N310 | S329 | I292 |
| 8x48 | K311 | P330 | R293 |
| 8x49 | Q312 | D331 | E294 |

**Table S3: Protein sequences of engineered receptors.**

| Name | Sequence |
|---|---|
| Opto-$\beta_2$AR-2.0<br><br>(Opto-$\beta_2$AR-All-In) | MNGTEGPNFYVPFSNKTGVVRSPFEAPQYYLAEPWQFSMLAAYMFLLIMLGFPINFLTLYVTVQHKKL<br>RTPLNYILLNLAVADLFMVFGGFTTTLYTSLHGYFVFGPTGCNLEGFFATLGGEIALWSLVVLAIERY<br>VAITSPFKYQSLLTENHAIMGVAFTWVMALACAAPPLVGWSRYIPEGMQCSCGIDYYTPHEETNNESF<br>VIYMFVVHFIIPLIVIFFCYGRVFQVAKRQLQKIDKSEGRFHSPNLGQVEQDGRSGHGLRRSSKFCLK<br>EHKALKTLIIMVIAFLICWLPYAGVAFYIFTHQGSDFGPIFMTIPAFFAKTSAVYNPVIYIMRSPDFR<br>IAFQELLCLRRSSSKAYGNGYSSNSNGKTDYMGEASGCQLGQEKESERLCEDPPGTESFVNCQGTVPS<br>LSLDSQGRNCSTNDSPL |
| Opto-$\beta_2$AR-Loops | MNGTEGPNFYVPFSNKTGVVRSPFEAPQYYLAEPWQFSMLAAYMFLLIMLGFPINFLTLYVTVQHKKL<br>RTPLNYILLNLAVADLFMVFGGFTTTLYTSLHGYFVFGPTGCNLEGFFATLGGEIALWSLVVLAIERY<br>VVVCSPFKYQSLLTENHAIMGVAFTWVMALACAAPPLVGWSRYIPEGMQCSCGIDYYTPHEETNNESF<br>VIYMFVVHFIIPLIVIFFCYGQLVFTVKEAAAQQQEGRFHSPNLGQVEQDGRSGHGLRRSSKFCATTQ<br>KAEKEVTRMVIIMVIAFLICWLPYAGVAFYIFTHQGSDFGPIFMTIPAFFAKTSAVYNPVIYIMMNKQ<br>FRIAFQELLCLRRSSSKAYGNGYSSNSNGKTDYMGEASGCQLGQEKESERLCEDPPGTESFVNCQGTV<br>PSLSLDSQGRNCSTNDSPL |
| Opto-$\beta_2$AR-2.0-Blue<br><br>(T118A-E122D-A318S) | MNGTEGPNFYVPFSNKTGVVRSPFEAPQYYLAEPWQFSMLAAYMFLLIMLGFPINFLTLYVTVQHKKL<br>RTPLNYILLNLAVADLFMVFGGFTTTLYTSLHGYFVFGPTGCNLEGFFAALGGDIALWSLVVLAIERY<br>VAITSPFKYQSLLTENHAIMGVAFTWVMALACAAPPLVGWSRYIPEGMQCSCGIDYYTPHEETNNESF<br>VIYMFVVHFIIPLIVIFFCYGRVFQVAKRQLQKIDKSEGRFHSPNLGQVEQDGRSGHGLRRSSKFCLK<br>EHKALKTLIIMVIAFLICWLPYAGVAFYIFTHQGSDFGPIFMTIPSFFAKTSAVYNPVIYIMRSPDFR<br>IAFQELLCLRRSSSKAYGNGYSSNSNGKTDYMGEASGCQLGQEKESERLCEDPPGTESFVNCQGTVPS<br>LSLDSQGRNCSTNDSPL |
| Opto-$\beta_2$AR-Blue<br><br>(T149A-E153D-A349S) | MKTIIALSYIFCLVFAMYTDIEMNRLGKDSLMNGTEGPNFYVPFSNKTGVVRSPFEAPQYYLAEPWQF<br>SMLAAYMFLLIMLGFPINFLTLYVIAKFERLQTVLNYILLNLAVADLFMVFGGFTTTLYTSLHGYFVF<br>GPTGCNLEGFFAALGGDIALWSLVVLAIERYVVVTSPFKYQSLLTKNKAIMGVAFTWVMALACAAPPL<br>VGWSRYIPEGMQCSCGIDYYTPHEETNNESFVIYMFVVHFIIPLIVIFFCYGRVFQVAKRQLQKIDKS<br>EGRFHSPNLGQVEQDGRSGHGLRRSSKFCLKEHKALRMVIIMVIAFLICWLPYAGVAFYIFTHQGSDF<br>GPIFMTIPSFFAKTSAVYNPVIYIMMNKQFRIAFQELLCLRRSSSKAYGNGYSSNSNGKTDYMGEASG<br>CQLGQEKESERLCEDPPGTESFVNCQGTVPSLSLDSQGRNCSTNDSPLTETSQVAPA |
| Opto-$A_{2A}$R-2.0 | MNGTEGPNFYVPFSNKTGVVRSPFEAPQYYLAEPWQFSMLAAYMFLLIMLGFPINFLTLYVTVQHKKL<br>RTPLNYILLNLAVADLFMVFGGFTTTLYTSLHGYFVFGPTGCNLEGFFATLGGEIALWSLVVLAIERY<br>VAIRIPLRYNGLVTENHAIMGVAFTWVMALACAAPPLVGWSRYIPEGMQCSCGIDYYTPHEETNNESF<br>VIYMFVVHFIIPLIVIFFCYGQIFLAARRQLKQMESQPLPGERARSTLQKEVHAAKSLIIMVIAFLIC<br>WLPYAGVAFYIFTHQGSDFGPIFMTIPAFFAKTSAVYNPVIYIMRIREFRQTFRKIIRSHVLRQQEPF<br>KAAGTSARILAAHGSDGEQVSLRLNGHPPGVWANGSAPHPERRPNGYALGLVSGGSAQESQGNTGLPD<br>VELLSHELKGVCPEPPGLDDPLAQDGAGVS |

#### Table S4: DNA sequences of engineered receptors.

| Name | Sequence |
|---|---|
| Opto-$\beta_2$AR-2.0<br><br>(Opto-$\beta_2$AR-All-In) | ATGAACGGGACCGAGGGCCCAAACTTCTACGTGCCTTTCTCCAACAAGACGGGCGTGGTGCGCAGCCCCT<br>TCGAGGCCCCGCAGTACTACCTGGCGGAGCCATGGCAGTTCTCCATGCTGGCCGCCTACATGTTCCTGCT<br>GATCATGCTTGGCTTCCCCATCAACTTCCTCACGCTGTACGTCACCGTGCAGCATAAAAAACTGCGCACC<br>CCGCTCAACTACATCCTGCTCAACCTGGCCGTGGCCGACCTCTTCATGGTGTTCGGGGGCTTCACCACCA<br>CCCTCTACACCTCTCTGCACGGGTACTTCGTCTTTGGGCCCACGGGCTGCAACCTGGAGGGCTTCTTTGC<br>CACCTTGGGCGGTGAAATTGCACTGTGGTCCTTGGTGGTCCTGGCCATCGAGCGGTACGTGGCGATTACA<br>TCGCCATTCAAGTACCAGAGCCTGCTGACCGAAAACCATGCCATCATGGGCGTCGCCTTCACCTGGGTCA<br>TGGCTCTGGCCTGTGCCGCGCCCCCCCTCGTCGGCTGGTCCAGGTACATCCCGGAGGGCATGCAGTGCTC<br>GTGCGGGATTGACTACTACACGCCCCACGAGGAAACCAACAATGAGTCGTTCGTCATCTACATGTTCGTG<br>GTCCACTTCATCATCCCCCTGATTGTCATATTCTTCTGCTACGGAAGGGTGTTCCAGGTGGCCAAAAGGC<br>AGCTCCAGAAGATAGACAAATCTGAGGGAAGATTCCACTCCCCAAACCTCGGCCAGGTGGAGCAGGATGG<br>GCGGAGTGGGCACGGACTCCGAAGGTCCTCCAAGTTCTGCTTGAAGGAGCACAAAGCCCTCAAAACCCTG<br>ATCATCATGGTCATCGCTTTCCTAATCTGCTGGCTGCCCTACGCTGGGGTGGCGTTCTACATCTTCACCC<br>ATCAGGGCTCTGACTTTGGCCCCATCTTCATGACCATCCCGGCTTTCTTTGCCAAGACTTCTGCCGTCTA<br>CAACCCCGTCATCTACATCATGCGCAGCCCGGATTTTCGCATTGCCTTCCAGGAGCTTCTATGCCTCCGC<br>AGGTCCTCTTCAAAAGCCTATGGGAACGGCTACTCCAGCAACAGTAATGGCAAAACAGACTACATGGGGG<br>AGGCGAGTGGATGTCAGCTGGGGCAGGAAAAAGAAAGTGAACGGCTGTGTGAGGACCCCCCAGGCACGGA<br>AAGCTTTGTGAACTGTCAAGGTACTGTGCCTAGCCTTAGCCTTGATTCCCAAGGGAGGAACTGTAGTACA<br>AATGACTCACCGCTGTAA |
| Opto-$\beta_2$AR-Loops | ATGAACGGGACCGAGGGCCCAAACTTCTACGTGCCTTTCTCCAACAAGACGGGCGTGGTGCGCAGCCCCT<br>TCGAGGCCCCGCAGTACTACCTGGCGGAGCCATGGCAGTTCTCCATGCTGGCCGCCTACATGTTCCTGCT<br>GATCATGCTTGGCTTCCCCATCAACTTCCTCACGCTGTACGTCACCGTGCAGCATAAAAAACTGCGCACC<br>CCGCTCAACTACATCCTGCTCAACCTGGCCGTGGCCGACCTCTTCATGGTGTTCGGGGGCTTCACCACCA<br>CCCTCTACACCTCTCTGCACGGGTACTTCGTCTTTGGGCCCACGGGCTGCAACCTGGAGGGCTTCTTTGC<br>CACCTTGGGCGGTGAAATTGCACTGTGGTCCTTGGTGGTCCTGGCCATCGAGCGGTACGTGGTGGTATGC<br>TCGCCATTCAAGTACCAGAGCCTGCTGACCGAAAACCATGCCATCATGGGCGTCGCCTTCACCTGGGTCA<br>TGGCTCTGGCCTGTGCCGCGCCCCCCCTCGTCGGCTGGTCCAGGTACATCCCGGAGGGCATGCAGTGCTC<br>GTGCGGGATTGACTACTACACGCCCCACGAGGAAACCAACAATGAGTCGTTCGTCATCTACATGTTCGTG<br>GTCCACTTCATCATCCCCCTGATTGTCATATTCTTCTGCTACGGACAACTCGTCTTCACGGTAAAGGAAG<br>CCGCTGCCCAGCAGCAAGAGGGGCGATTCCATTCACCGAATTTGGGCCAGGTGGAACAAGACGGACGGTC<br>TGGGCATGGACTTAGACGATCTTCTAAATTCTGTGCTACTACTCAAAAGGCTGAGAAGGAGGTTACAAGG<br>ATGGTAATCATCATGGTCATCGCTTTCCTAATCTGCTGGCTGCCCTACGCTGGGGTGGCGTTCTACATCT<br>TCACCCATCAGGGCTCTGACTTTGGCCCCATCTTCATGACCATCCCGGCTTTCTTTGCCAAGACTTCTGC<br>CGTCTACAACCCCGTCATCTACATCATGATGAACAAGCAGTTCCGCATTGCCTTCCAGGAGCTTCTATGC<br>CTCCGCAGGTCCTCTTCAAAAGCCTATGGGAACGGCTACTCCAGCAACAGTAATGGCAAAACAGACTACA<br>TGGGGGAGGCGAGTGGATGTCAGCTGGGGCAGGAAAAAGAAAGTGAACGGCTGTGTGAGGACCCCCCAGG<br>CACGGAAAGCTTTGTGAACTGTCAAGGTACTGTGCCTAGCCTTAGCCTTGATTCCCAAGGGAGGAACTGT<br>AGTACAAATGACTCACCGCTGTAA |
| Opto-$\beta_2$AR-2.0-Blue<br><br>(T118A-E122D-A318S) | ATGAACGGGACCGAGGGCCCAAACTTCTACGTGCCTTTCTCCAACAAGACGGGCGTGGTGCGCAGCCCCT<br>TCGAGGCCCCGCAGTACTACCTGGCGGAGCCATGGCAGTTCTCCATGCTGGCCGCCTACATGTTCCTGCT<br>GATCATGCTTGGCTTCCCCATCAACTTCCTCACGCTGTACGTCACCGTGCAGCATAAAAAACTGCGCACC<br>CCGCTCAACTACATCCTGCTCAACCTGGCCGTGGCCGACCTCTTCATGGTGTTCGGGGGCTTCACCACCA<br>CCCTCTACACCTCTCTGCACGGGTACTTCGTCTTTGGGCCCACGGGCTGCAACCTGGAGGGCTTCTTTGC<br>CgCCTTGGGCGGTGAtATTGCACTGTGGTCCTTGGTGGTCCTGGCCATCGAGCGGTACGTGGCGATTACA<br>TCGCCATTCAAGTACCAGAGCCTGCTGACCGAAAACCATGCCATCATGGGCGTCGCCTTCACCTGGGTCA |

| | |
|---|---|
| | TGGCTCTGGCCTGTGCCGCGCCCCCCCTCGTCGGCTGGTCCAGGTACATCCCGGAGGGCATGCAGTGCTC<br>GTGCGGGATTGACTACTACACGCCCCACGAGGAAACCAACAATGAGTCGTTCGTCATCTACATGTTCGTG<br>GTCCACTTCATCATCCCCCTGATTGTCATATTCTTCTGCTACGGAAGGGTGTTCCAGGTGGCCAAAAGGC<br>AGCTCCAGAAGATAGACAAATCTGAGGGAAGATTCCACTCCCCAAACCTCGGCCAGGTGGAGCAGGATGG<br>GCGGAGTGGGCACGGACTCCGAAGGTCCTCCAAGTTCTGCTTGAAGGAGCACAAAGCCCTCAAAACCCTG<br>ATCATCATGGTCATCGCTTTCCTAATCTGCTGGCTGCCCTACGCTGGGGTGGCGTTCTACATCTTCACCC<br>ATCAGGGCTCTGACTTTGGCCCCATCTTCATGACCATCCCGtCTTTCTTTGCCAAGACTTCTGCCGTCTA<br>CAACCCCGTCATCTACATCATGCGCAGCCCGGATTTTCGCATTGCCTTCCAGGAGCTTCTATGCCTCCGC<br>AGGTCCTCTTCAAAAGCCTATGGGAACGGCTACTCCAGCAACAGTAATGGCAAAACAGACTACATGGGGG<br>AGGCGAGTGGATGTCAGCTGGGGCAGGAAAAAGAAAGTGAACGGCTGTGTGAGGACCCCCCAGGCACGGA<br>AAGCTTTGTGAACTGTCAAGGTACTGTGCCTAGCCTTAGCCTTGATTCCCAAGGGAGGAACTGTAGTACA<br>AATGACTCACCGCTGTAA |
| Opto-$\beta_2$AR-Blue<br><br>(T149A-E153D-A349S) | ATGAAAACGATCATCGCCCTGAGCTACATCTTCTGCCTGGTATTCGCCATGTACACCGATATAGAGATGA<br>ACAGGCTGGGAAAGGATAGCCTCATGAACGGGACCGAGGGCCCAAACTTCTACGTGCCTTTCTCCAACAA<br>GACGGGCGTGGTGCGCAGCCCCTTCGAGGCCCCGCAGTACTACCTGGCGGAGCCATGGCAGTTCTCCATG<br>CTGGCCGCCTACATGTTCCTGCTGATCATGCTTGGCTTCCCCATCAACTTCCTCACGCTGTACGTCATTG<br>CCAAGTTCGAGAGGCTACAGACTGTCCTCAACTACATCCTGCTCAACCTGGCCGTGGCCGACCTCTTCAT<br>GGTGTTCGGGGGCTTCACCACCACCCTCTACACCTCTCTGCACGGGTACTTCGTCTTTGGGCCCACGGGC<br>TGCAACCTGGAGGGCTTCTTTGCCGCCTTGGGCGGTGATATTGCACTGTGGTCCTTGGTGGTCCTGGCCA<br>TCGAGCGGTACGTGGTGGTGACATCGCCATTCAAGTACCAGAGCCTGCTGACCAAGAATAAGGCCATCAT<br>GGGCGTCGCCTTCACCTGGGTCATGGCTCTGGCCTGTGCCGCGCCCCCCCTCGTCGGCTGGTCCAGGTAC<br>ATCCCGGAGGGCATGCAGTGCTCGTGCGGGATTGACTACTACACGCCCCACGAGGAAACCAACAATGAGT<br>CGTTCGTCATCTACATGTTCGTGGTCCACTTCATCATCCCCCTGATTGTCATATTCTTCTGCTACGGAAG<br>GGTGTTCCAGGTGGCCAAAAGGCAGCTCCAGAAGATAGACAAATCTGAGGGAAGATTCCACTCCCCAAAC<br>CTCGGCCAGGTGGAGCAGGATGGGCGGAGTGGGCACGGACTCCGAAGGTCCTCCAAGTTCTGCTTGAAGG<br>AGCACAAAGCCCTCCGCATGGTGATCATCATGGTCATCGCTTTCCTAATCTGCTGGCTGCCCTACGCTGG<br>GGTGGCGTTCTACATCTTCACCCATCAGGGCTCTGACTTTGGCCCCATCTTCATGACCATCCCGTCTTTC<br>TTTGCCAAGACTTCTGCCGTCTACAACCCCGTCATCTACATCATGATGAACAAGCAGTTCCGCATTGCCT<br>TCCAGGAGCTTCTATGCCTCCGCAGGTCCTCTTCAAAAGCCTATGGGAACGGCTACTCCAGCAACAGTAA<br>TGGCAAAACAGACTACATGGGGGAGGCGAGTGGATGTCAGCTGGGGCAGGAAAAAGAAAGTGAACGGCTG<br>TGTGAGGACCCCCCAGGCACGGAAAGCTTTGTGAACTGTCAAGGTACTGTGCCTAGCCTTAGCCTTGATT<br>CCCAAGGGAGGAACTGTAGTACAAATGACTCACCGCTGACAGAAACCAGCCAAGTGGCGCCTGCCTAA |
| Opto-$A_{2A}$R-2.0 | ATGAACGGGACCGAGGGCCCAAACTTCTACGTGCCTTTCTCCAACAAGACGGGCGTGGTGCGCAGCCCCT<br>TCGAGGCCCCGCAGTACTACCTGGCGGAGCCATGGCAGTTCTCCATGCTGGCCGCCTACATGTTCCTGCT<br>GATCATGCTTGGCTTCCCCATCAACTTCCTCACGCTGTACGTCACCGTGCAGCATAAAAAACTGCGCACC<br>CCGCTCAACTACATCCTGCTCAACCTGGCCGTGGCCGACCTCTTCATGGTGTTCGGGGGCTTCACCACCA<br>CCCTCTACACCTCTCTGCACGGGTACTTCGTCTTTGGGCCCACGGGCTGCAACCTGGAAGGCTTCTTTGC<br>CACCTTGGGCGGTGAAATTGCACTGTGGTCCTTGGTGGTCCTGGCCATCGAGCGGTACGTGGCGATTAGG<br>ATACCGCTGCGGTATAACGGGCTTGTTACGGAAAACCATGCCATCATGGGCGTCGCCTTCACCTGGGTCA<br>TGGCTCTGGCCTGTGCCGCGCCCCCCCTCGTCGGCTGGTCCAGGTACATCCCGGAGGGCATGCAGTGCTC<br>GTGCGGGATTGACTACTACACGCCCCACGAGGAAACCAACAATGAGTCGTTCGTCATCTACATGTTCGTG<br>GTCCACTTCATCATCCCCCTGATTGTCATATTCTTCTGCTACGGACAGATATTCCTGGCGGCAAGACGCC<br>AACTGAAGCAGATGGAAAGCCAACCGTTGCCCGGGGAACGCGCGCGGAGCACCCTTCAAAAAGAAGTCCA<br>TGCAGCCAAGAGTCTCATCATCATGGTCATCGCTTTCCTAATCTGCTGGCTGCCCTACGCTGGGGTGGCG<br>TTCTACATCTTCACCCATCAGGGCTCTGACTTTGGCCCCATCTTCATGACCATCCCGGCTTTCTTTGCCA<br>AGACTTCTGCCGTCTACAACCCCGTCATCTACATCATGCGGATTCGGGAGTTCAGACAGACCTTTAGAAA<br>AATTATAAGATCTCACGTCCTGAGGCAACAAGAACCCTTTAAAGCTGCGGGGACTTCTGCCCGGATTTTG<br>GCGGCGCATGGGAGCGATGGTGAACAGGTGTCCCTCAGATTGAACGGGCATCCTCCGGGGGTGTGGGCTA |

| | |
|---|---|
| | ATGGTAGTGCTCCTCACCCTGAAAGACGACCTAACGGGTACGCACTGGGACTTGTGTCAGGTGGGTCTGC<br>CCAGGAAAGCCAAGGTAACACGGGGCTTCCTGACGTAGAGTTGTTGTCTCATGAGCTGAAGGGTGTCTGC<br>CCTGAACCTCCTGGTCTGGACGACCCGCTCGCACAGGATGGAGCAGGTGTTAGTTAA |
